# Structural and Mechanistic Basis for Antibody Neutralization of the Measles Fusion Protein

**DOI:** 10.1101/2025.09.11.675466

**Authors:** Dawid S. Zyla, Roberta Della Marca, Davide Lacarbonara, Gele Niemeyer, Gillian Zipursky, Laura Di Clemente, Gabriella Jonathan-Trakht, Gavreel Kalantarov, Marissa Acciani, Giulia Laterza, Dariia Vyshenska, Kathryn M Hastie, Branka Horvat, Alexander L. Greninger, Stefan Niewiesk, Erica Ollmann Saphire, Matteo Porotto

**Author notes:** These authors contributed equally (Dawid S. Zyla, Roberta Della Marca).

## Abstract

Measles virus (MeV) is a highly contagious viral pathogen and remains a major global health threat. Resurgent infections, driven by insufficient vaccine coverage, waning herd immunity, and the vulnerability of immunocompromised individuals, highlight the urgent need for effective countermeasures. Monoclonal antibodies (mAbs) represent a promising strategy, both as antiviral agents and as probes of viral entry mechanisms. While most vaccine-elicited neutralizing antibodies target the hemagglutinin (H) protein, emerging evidence suggests that antibodies against the fusion (F) protein are also potent inhibitors. Still, there is insufficient information on the target sites and activities of antibodies against the F protein. Like other class I fusion proteins, MeV F exists in a metastable prefusion state that undergoes dramatic conformational changes during viral entry.

Here, we selected four mAbs that recognize conformational patterns of F-prefusion and/or postfusion, characterized their epitopes, specificities, and antiviral activities. Structural analyses mapped antibody interactions onto pre- and postfusion F conformations, revealing that all three neutralizing mAbs are specific for the prefusion form, while the non-neutralizing mAb recognizes only the postfusion F. Biophysical and functional assays defined distinct mechanisms: neutralization occurs either by stabilizing the prefusion protein or by preventing the extended intermediate from completing fusion. We also describe a novel mechanism of neutralization in which an antibody prematurely triggers F activation but blocks the subsequent refolding required for viral entry. Together, these findings provide the first detailed mapping of neutralizing epitopes on the MeV F protein and establish a framework for the rational design of F-targeted intervention.

## Introduction

Measles is a highly contagious viral disease that has reemerged globally in recent years. Achieving widespread vaccine-induced immunity is critical to preventing outbreaks, particularly as unvaccinated and immunocompromised individuals remain at high risk for severe disease. Although the measles vaccine has an excellent long-term safety and efficacy record, neither global nor American vaccination rates have reached the 95% threshold required for herd immunity^1–6^. Consequently, the recent resurgence in measles cases reflects a failure to vaccinate rather than a failure of the vaccine itself. Over the past five years, declining vaccination rates have been fueled by disruptions to routine immunization programs during the COVID-19 pandemic and by the increasing impact of vaccine hesitancy^1–6^.

These developments increase the vulnerability of individuals who cannot be vaccinated due to medical contraindications. Post-exposure prophylaxis (PEP) with total immune globulin remains the only recommended option for these individuals; however, its reliability is threatened by declining measles antibody titers within the donor pool^1,7–15^. With measles elimination efforts stalled and a growing number of high-risk individuals, there is an urgent need for effective antiviral therapies. However, currently, there is no antiviral treatment yet approved for measles. Passive immunization using monoclonal or polyclonal antibodies has long been a cornerstone in preventing and treating other viral infections, particularly in individuals who are unable to receive active vaccination. However, only a limited number of monoclonal antibodies (mAbs) have received clinical approval for the treatment of those viral infectious diseases^16–18^, and none are available for measles.

Antibodies can exert antiviral effects by targeting viral glycoproteins critical for host cell entry. In measles virus (MeV), this process involves two envelope proteins: the hemagglutinin (H) protein, which binds host receptors, and the fusion (F) protein, which mediates membrane fusion. Fusion is a dynamic, multistep process: following H–F interaction and receptor engagement, F transitions through an extended “prehairpin” intermediate, during which its heptad repeat regions (HRN and HRC) span the viral and host membranes. These regions then fold into a six-helix bundle (6HB), completing fusion and enabling viral entry.

MAbs targeting MeV H and F offer several advantages: they can neutralize the virus early in infection, avoid the toxicity and off-target effects of small molecules, and provide protection for immunocompromised or vaccine-ineligible individuals^1,16^. Historical studies suggest that vaccine-induced antibodies primarily target the H protein, but natural infection elicits robust neutralizing antibodies against both H and F, indicating that F is a promising target for mAbs development^19,20^. Strategies to block viral entry include targeting: (i) H protein binding to hSLAM/Nectin-4 receptors, (ii) F activation via H–F interaction, (iii) conformational transitions exposing the F fusion peptide, and (iv) F refolding and pore expansion. MeV F is a class I trimeric fusion protein undergoing substantial conformational rearrangements from prefusion to postfusion states, a feature shared with many other enveloped viruses^21–24^. We have generated and characterized a panel of chimeric (mouse Fab/human Fc) mAbs targeting MeV F and present here four antibodies selected for distinct epitope specificity and mechanism of action. Three of these mAbs exhibit potent neutralization against the two currently circulating wild-type MeV strains, providing a strong foundation for antiviral development. Our data demonstrate that the MeV F protein contains multiple neutralization-sensitive epitopes that can be targeted by potent mAbs. By combining antibodies with distinct mechanisms, broad and synergistic inhibition can be achieved, raising the barrier to resistance. A cocktail of long-acting humanized anti-F antibodies could serve as a critical countermeasure for individuals unable to receive vaccination, for outbreak control, and as a complement to existing PEP strategies.

## Results

### Neutralization Activity and Fusion Protein Reactivity of Chimeric Monoclonal Antibodies Against Measles Virus

We generated and characterized a panel of mouse monoclonal antibodies (mAbs) targeting the MeV F protein. The Fab regions of four of these antibodies were grafted onto a common human Fc backbone. Here, we present the functional and structural properties of these four chimeric mAbs, termed 77.1, Y10F, H8, and C6.

We first assessed their ability to recognize F glycoproteins expressed in HEK293T cells from two currently circulating wild-type strains (B3 and D8), the vaccine strain (Moraten), and two previously characterized B3 F variants – L454W, which destabilizes the prefusion conformation^25,26^, and E455G, which stabilizes the prefusion conformation^26,27^ (**Fig. 1A**). MAbs 77.1 and Y10F recognized each of currently circulating wild-type F proteins, the Moraten-vaccine F, and the prefusion-stabilized E455G variant. MAb 77.1 and Y10F recognized the B3 F carrying the destabilizing L454W mutation; however, the recognition was markedly reduced. In contrast, mAb H8 recognized all F proteins tested, while mAb C6 did not recognize wild-type or stabilized E455G F variant and recognized only the Moraten-vaccine F and the destabilized L454W variant^25^.

**Figure 1.**
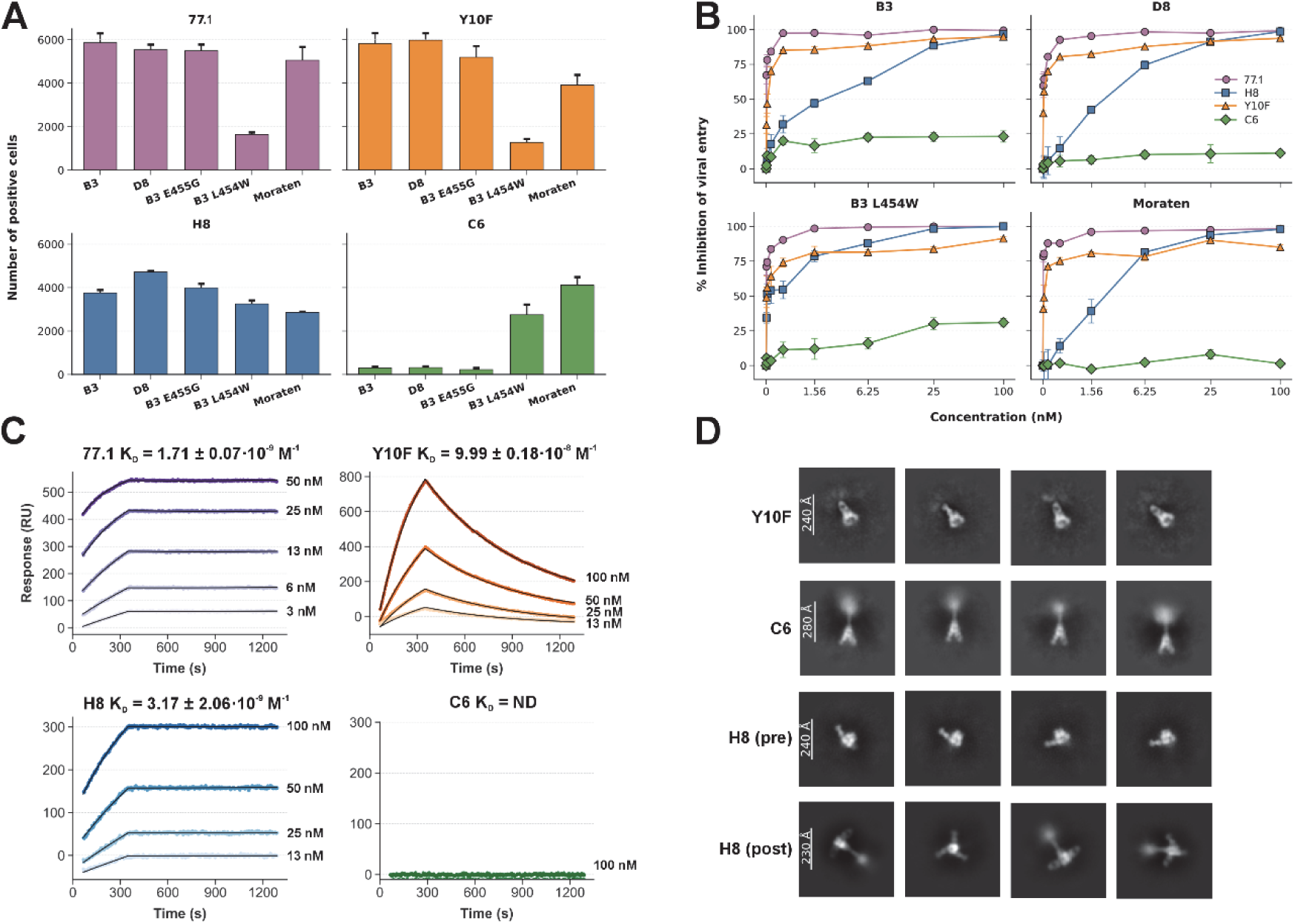
Epitope Recognition and Neutralizing Activity of Anti-F Chimeric Monoclonal Antibodies. A) Recognition of cell-surfaced expressed fusion protein from the wild-type B3 and D8 strains (the only two currently circulating strains), vaccine strain Moraten, destabilized mutant B3 L454W, and stabilized mutant B3 E455G. Cells expressing F variants were incubated for 24 hours and analyzed for binding to chimeric monoclonal antibodies (Y10F, 77.1, H8, and C6). Error bars represent the mean ± SEM, and bar heights represent the mean of the measurement. Data are from at least 3 separate experiments. B) Antiviral potency of mAbs measured by plaque reduction neutralization tests (PRNT). We assessed the antiviral potency of the four mAbs in a viral entry assay using two circulating wild-type strains (B3 and D8), a vaccine strain (Moraten), and a neuropathogenic variant (B3 L454W F). Vero-hSLAM/CD150 cells were infected with approximately 250 plaque-forming units (PFU)/well of each virus in the presence of increasing concentrations of mAbs 77.1 (violet circles), H8 (blue squares), Y10F (red triangles), and C6 (green diamonds) as indicated on the x-axis. After 2 hours, the inoculum was removed and replaced with 1% methylcellulose. Infection was monitored for 72 hours, with images acquired every 24 hours using a Cytation 5 imager. Plaques were quantified at 48 hours post-infection. Data represent mean ± SEM from three independent experiments. C) Surface plasmon resonance (SPR) sensorgrams collected on a Carterra LSA with immobilized antibodies and prefusion F_ECTO_ as the analyte. Association and dissociation phases were globally fit to a 1:1 Langmuir model across a concentration series to obtain k_on_ and k_off_; equilibrium dissociation constants (K_D_=k_off_/k_on_) are reported for each antibody. D) 2D class averages obtained from negative-stain electron microscopy (NS-EM) of the Fabs-F_ECTO_ complexes.

Neutralization potency, measured by plaque reduction neutralization tests (PRNT) against four recombinant viruses (B3, D8, B3 F-L454W, and vaccine strain Moraten), revealed that 77.1 and Y10F were the most potent antibodies, with IC_90_ values ranging from 0.15–0.5 nM for 77.1 and 7–83 nM for Y10F. H8 was less potent overall (IC_90_: 12–64 nM), but showed significantly higher activity against the neuropathogenic F-destabilized B3 F-L454W^28^ variant (20 nM) compared to Y10F (83 nM). In contrast, C6 exhibited no measurable neutralizing activity.

We next quantified antibody binding to purified, stabilized, soluble prefusion F ectodomain (F_ECTO_^27^) using high-throughput surface plasmon resonance (SPR) on a Carterra LSA (**Fig. 1C)**. This F contains natural substitutions in place of disulfide bridges, which stabilize the prefusion state while still permitting conformational transition, albeit in a more gradual and controllable manner than wild-type F^27^. Three antibodies (77.1, H8, Y10F) bound this purified, prefusion measles F, while the postfusion specific control mAb C6 showed no detectable binding to this prefusion F, as expected for its postfusion conformation-specific recognition.

We observed distinct kinetic signatures across the three binders. mAb 77.1 showed the highest affinity (K_D_ = 1.71 ± 0.01 × 10⁻⁹ M). H8 had the slowest association and the strongest dissociation, with a KD of 3.1 ± 2.0 × 10⁻⁹ M. Y10F associated faster than H8 and 77.1, but dissociated more quickly, yielding a lower overall affinity (KD = 9.99 ± 0.18 × 10⁻⁸ M). These trends indicate that Y10F reaches its site rapidly but forms a shorter-lived complex (**Fig.1C** and **Table 1**).

**Table 1.** Fitting parameters of antibody kinetics data from LSA.

| Antibody | H8 | Y10F | 77.1 | C6 |
| --- | --- | --- | --- | --- |
| $k_a$ (M<sup>-1</sup>s<sup>-1</sup>) | 23640 ± 810 | 17900 ± 300 | 121000 ± 2000 | ND |
| $k_d$ (s<sup>-1</sup>) | (7.48 ± 4.82) × 10<sup>-5</sup> | (1.790 ± 0.009) ×<br>10<sup>-3</sup> | (2.07 ± 0.09) × 10<sup>-4</sup> | ND |
| $K_D$ (M) | (3.17 ± 2.06) × 10<sup>-9</sup> | (9.99 ± 0.18) × 10<sup>-8</sup> | (1.71 ± 0.07) × 10<sup>-9</sup> | ND |
| $R^2$ | 0.99967 | 0.99942 | 0.99987 | ND |

ND = Not determined

To visualize the conformational preferences of each antibody, we performed negative-stain EM (NS-EM) on Fab–F_ECTO_ complexes (**Fig. 1D**). Class averages showed that Y10F engages the prefusion apex of F, whereas H8 can bind both prefusion and postfusion conformations. In contrast, C6 produced discernible complexes only with particles that had transitioned to the postfusion state. Taken together, these results indicate that three mAbs that bind prefusion epitopes are neutralizing (77.1, Y10F, H8), while the mAb that binds only a postfusion epitope is not neutralizing (C6).

### Stability of the measles fusion protein in antibody complexes

To assess how antibody binding affects the stability of the prefusion F_ECTO_, we performed nano differential scanning fluorimetry (nanoDSF; NanoTemper Prometheus Panta) (**Fig. 2A**). F_ECTO_ was incubated with a threefold molar excess of Fab for 30 min at room temperature. F_ECTO_ alone exhibited a melting temperature (Tm) of 49.7°C, consistent with prior measurements^27^. The presence of the postfusion-specific C6 did not affect the F_ECTO_ Tm. Although all three prefusion-specific antibodies (77.1, Y10F, and H8) altered F_ECTO_ thermal stability, they did so in distinct ways. Specifically, 77.1 increased the F_ECTO_ Tm by 3.8°C to 53.5°C, and Y10F increased it by 7.2°C to 56.9°C. In contrast, H8 decreased the F_ECTO_ Tm by 5.1°C to 44.6°C. To exclude contributions from Fab unfolding to these transitions, we recorded Fab melting profiles in the absence of F_ECTO_. The Fab-alone melting transitions matched the Fab-associated features observed in the complex traces, and there were no pronounced signal changes at the complex Tm attributable to Fab denaturation, indicating that the temperature shifts reported above reflect unfolding or conformational changes in F_ECTO_ (**Fig. 2B**).

**Figure 2.**
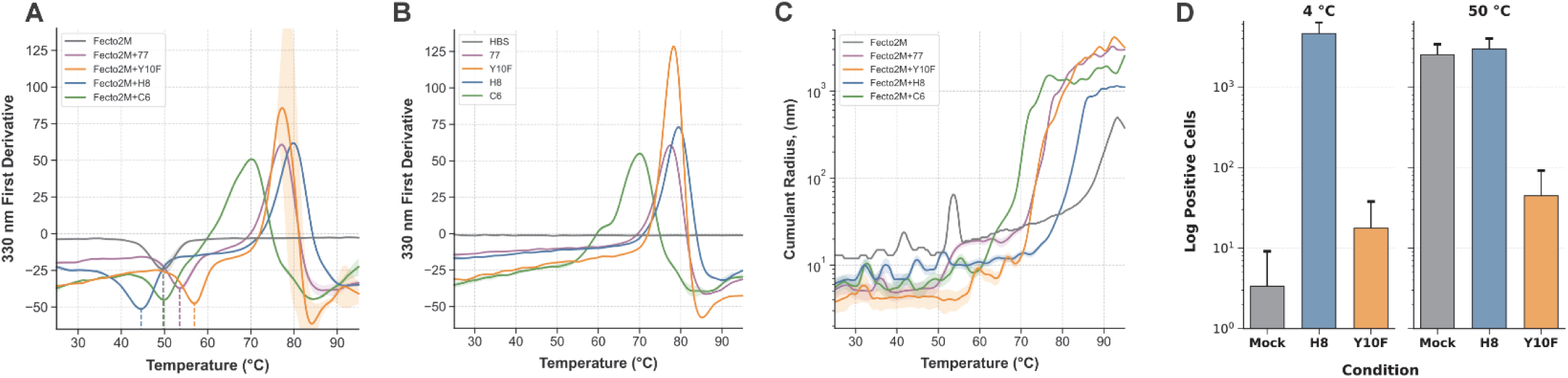
Thermal stability, size, and conformational effects of F-mAb complexes. A) F_ECTO_–Fab complexes (F_ECTO_–77, F_ECTO_–Y10F, F_ECTO_–H8, and F_ECTO_–C6) were assembled 30 min before measurement, loaded into capillaries, and monitored for thermal unfolding by intrinsic fluorescence at 330 nm. First-derivative nanoDSF traces for F_ECTO_–Fab complexes. Dashed, color-coded lines mark F_ECTO_ unfolding temperatures: F_ECTO_ alone, 49.7°C; with 77, 53.5°C; with Y10F, 56.9°C; with H8, 44.6°C; with C6, 49.9°C. B) First-derivative traces for Fab fragments alone; Fab melting temperatures were consistent with those in panel A: 77, 77.5°C; Y10F, 78.1°C; H8, 79.4°C; C6, 70.1°C. C) In parallel, dynamic light scattering (DLS) was recorded, and the cumulant radius was plotted as a function of temperature (°C). D) C6 binding to post-triggered F in the presence of H8 or Y10F, measured after incubation at 4°C or following a 50°C temperature shift. To assess the effect of H8 and Y10F on native full-length MeV F stability, cells expressing wild-type B3 F were incubated with 200 nM of either antibody. Exposure of the post-triggered F epitope was measured using C6 after 90 min at 4°C or following a temperature shift (30 min at 4°C then 30 min at 50°C). The percentage of C6-positive cells, reflecting post-triggered F exposure, is plotted on the y-axis. Error bars represent the mean ± SEM, and bar heights represent the mean of the measurement. Data are from at least 3 separate experiments.

In parallel, we monitored the cumulant (hydrodynamic) radius as a function of temperature (**Fig. 2C**). F_ECTO_ alone maintained a radius of ∼13 nm up to its Tm, after which the radius increased continuously, indicating aggregation. In contrast, all F_ECTO_–antibody mixtures maintained more compact radii (∼8–9 nm) over this range. For the F_ECTO_–77.1 complex, we observed stabilization of an intermediate state between ∼55°C and 70°C after triggering, consistent with our prior cryoEM structure demonstrating arrest at a fusion intermediate^27^. This intermediate state was followed by aggregation after the Fab melting temperature of 75°C. Other mAbs showed different results: the F_ECTO_–Y10F complex displayed a lower cumulant radius below the F_ECTO_ Tm and a uniform radius increase above, while the F_ECTO_–H8 complex displayed a rise in cumulant radius only above the Fab melting temperature. In contrast, in the presence of C6, aggregation began at ∼60°C, coinciding with the C6 Fab unfolding temperature. Given that C6 does not bind prefusion F_ECTO_, this behavior is consistent with the lack of interaction under these conditions. Together, the nanoDSF and size analyses demonstrate that the prefusion-specific mAbs 77.1, Y10F, and H8 exert strong but diverging effects on F_ECTO_ conformational stability, whereas C6 does not measurably affect the stability of the prefusion ectodomain.

Collectively, these biophysical analyses demonstrate that prefusion-specific mAbs 77.1, Y10F, and H8 strongly, but divergently, modulate F_ECTO_ conformational stability, whereas C6 has no measurable effect. Mechanistically, the data suggest opposing modes of action for Y10F and H8: Y10F stabilizes the prefusion conformation and raises the energy barrier for activation, thereby preventing fusion triggering, whereas H8 lowers the activation barrier and promotes premature triggering of the fusion machinery.

To validate these mechanistic insights in the context of native membrane-anchored F, we performed cell-surface staining with the postfusion-specific mAb C6 after preincubating cells at either 4°C or 50°C (the latter to thermally induce F triggering) in the presence of Y10F or H8 (**Fig. 2D**). Because C6 recognizes an epitope exposed only upon F triggering, its binding serves as a proxy for F activation. At 4°C, C6 bound F in the presence of H8, but not in the presence of Y10F. At 50°C, C6 failed to bind F in the presence of Y10F. These findings corroborate the biophysical data, supporting a model in which Y10F raises the conformational activation threshold of F, thereby maintaining the prefusion state, while H8 lowers this barrier and drives premature activation.

### Cryo-EM mapping of Y10F and H8 epitopes on the measles F protein

To define the antibody epitopes on F, we determined cryo-EM structures of the F_ECTO_ in complex with Y10F alone, with Y10F and 77.1 bound simultaneously, and with H8 alone.

The F_ECTO_–Y10F dataset exhibited mild preferred orientation, with many particles showing a tilted top view (**Supplemental Fig. 1**). Despite this, we reconstructed a 2.59 Å map that revealed asymmetric binding of a single Y10F Fab at the apex of F_ECTO_ (**Fig. 2A**). Y10F binds the region in the center of the apex, surrounded by the six glycosylation sites positioned on the three F2 subunits at positions 61 and 67 (**Fig. 3A**). Both heavy and light chains contribute to a dense interaction network. The heavy chain forms nine hydrogen bonds distributed across three protomers (including one F2 subunit and two F1 subunits), with residues N55 and N31 accounting for six of these nine bonds (**Table S1**). The light chain forms five hydrogen bonds with the same two F1 chains. The buried interface area is ∼870 Å², ∼70% contributed by the heavy chain, with an estimated by PISA binding free energy (ΔG) of −8.6 kcal/mol, ∼85% of which arises from heavy-chain contacts (**Fig. 3B**).

**Figure 3.**
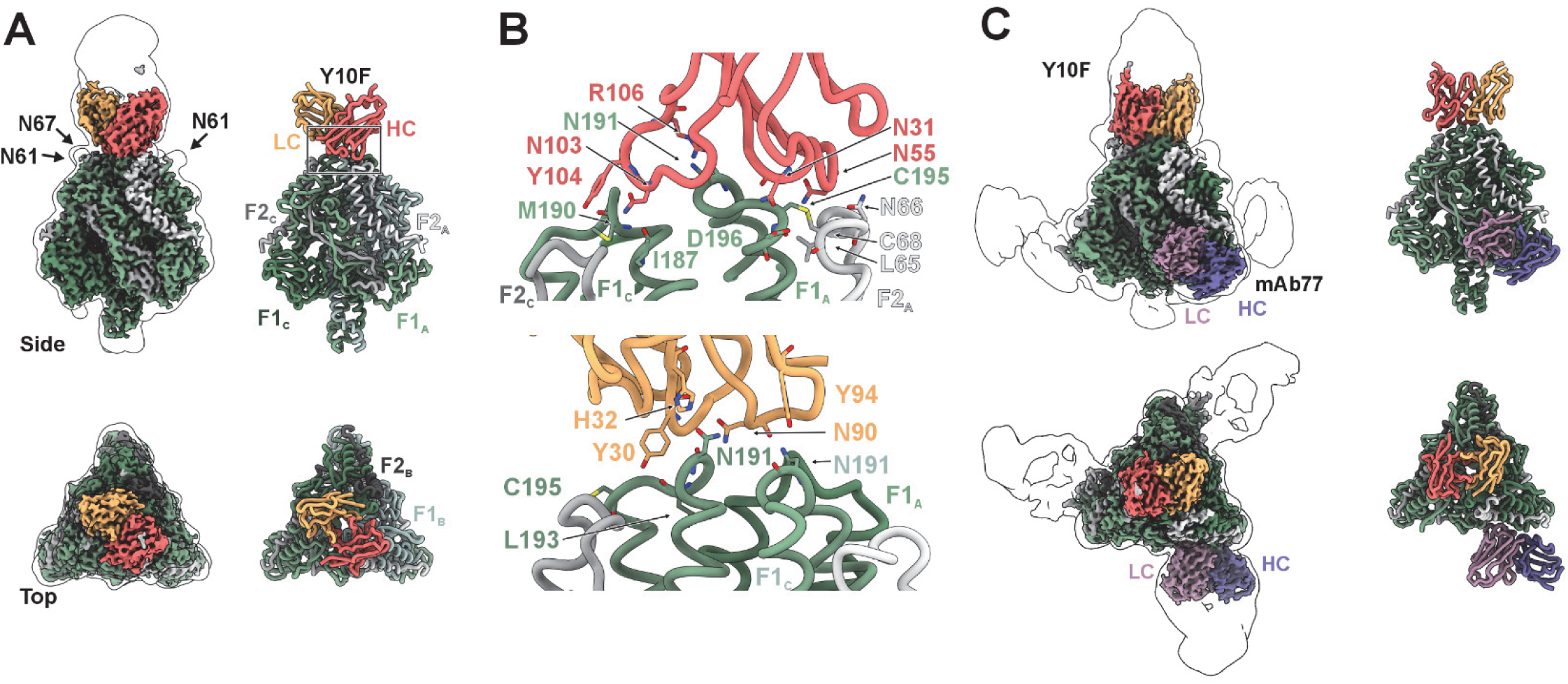
Cryo-EM analysis of F_ECTO_–Y10F and dual F_ECTO_–Y10F–77 complexes. **A)** Cryo-EM density and atomic model of the F_ECTO_ trimer with a single Y10F Fab bound asymmetrically at the apex. The model is shown as ribbons: Y10F heavy chain in light red, light chain in yellow; F1 protomers in shades of green and F2 protomers in shades of gray. Upper and lower panels show side and top views, respectively. Glycans near Y10F Fab are indicated with arrows. **B)** Epitope–paratope contacts between Y10F and F_ECTO_. The top panel highlights heavy-chain interactions, and the bottom panel highlights light-chain interactions. Both chains engage an asymmetric epitope on F. Interacting residues are color-coded by chain, matching the coloring from **A**). **C)** Cryo-EM map and model of F_ECTO_ bound by Y10F and 77 Fabs. Due to a symmetry mismatch (C1 for Y10F and C3 for 77) and partial Fab occupancy, only one copy of 77 is resolved in the final locally refined reconstruction.

We also solved the structure of F_ECTO_ bound simultaneously by Y10F and 77.1 (**Fig. 3C**) to test co-occupancy along with any binding-induced conformational changes that result from simultaneous binding of these two antibodies. Although both Fabs were supplied in excess, the complex displayed partial occupancy of each Fab, with individual particles displaying one, two, or three copies of 77.1. The combination of preferred orientation, asymmetric nature of the complex, and incomplete occupancy complicated particle selection. However, through symmetry expansion and focused masking, we obtained a reconstruction in which one copy each of 77.1 and Y10F were fully resolved. Superposition of the Y10F-only and Y10F+77.1 models yielded a Cα RMSD of 0.79 Å, indicating no significant global rearrangement of F_ECTO_. Local differences were restricted to the 77.1 epitope, particularly residues 433–442, which showed an RMSD of ∼2 Å, consistent with prior descriptions of 77.1-induced adjustments^27^. These data demonstrate that Y10F and 77.1 can co-bind F, supporting their use in an antibody cocktail.

To characterize H8 binding in both conformational states of F, we used two sample-preparation strategies guided by negative-stain EM (**Fig. 1D**) and nanoDSF (**Fig. 2**) results. To solve a prefusion complex (**Fig. 4A**), F_ECTO_ and H8 Fab were incubated for 15 min at room temperature before vitrification to promote complex formation while minimizing triggering to the postfusion state; longer incubations increased the proportion of postfusion particles. To solve a postfusion complex (**Fig. 4B**), F_ECTO_ and H8 were incubated overnight at 4°C and then applied to GO grids.

**Figure 4.**
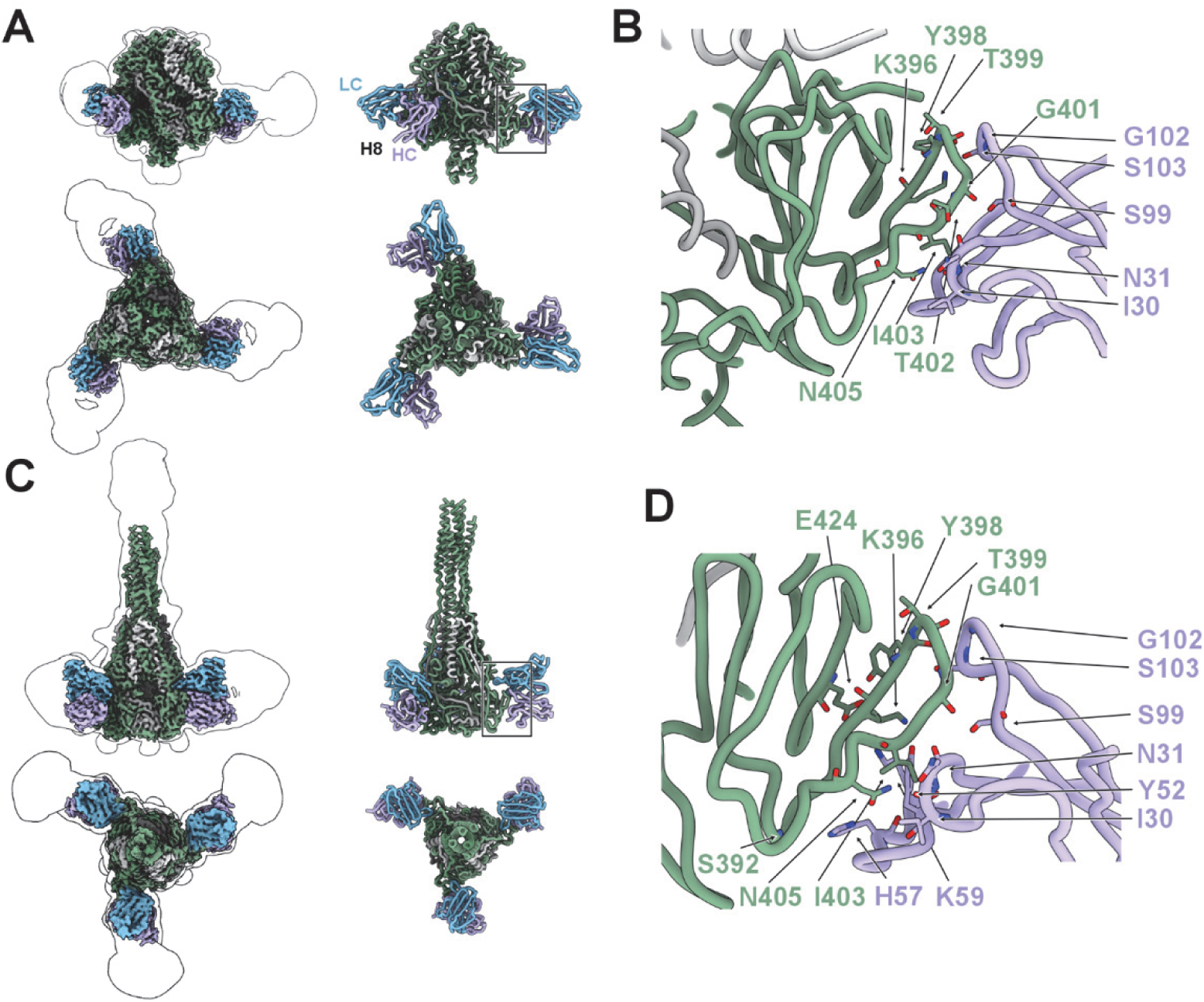
Cryo-EM structures of H8 bound to the measles F_ECTO_ in pre- and postfusion states. A) Cryo-EM density and atomic model of prefusion F_ECTO_ symmetrically bound by three copies of Fab H8. Ribbon representation shows H8 heavy chains in light purple and light chains in light blue. Side (top) and top (bottom) views are shown. B) Close-up of the prefusion interface. H8 engages the outer face of domain II exclusively through its heavy chain. C) Cryo-EM density and atomic model of postfusion F_ECTO_ with three copies of Fab H8 bound. Side (top) and top (bottom) views are shown. D) Close-up of the postfusion interface. As in the prefusion state, H8 binds exclusively via its heavy chain and targets the same region on domain II.

In the prefusion state, H8 bound symmetrically to all three protomers at an epitope on the outer face of domain II, opposite the F vertex contacted by 77.1, producing a triskelion-like structure. The epitope centered on the single loop in domain II (residues 396–405). Binding was dominated by the heavy chain and was stabilized by ten hydrogen bonds. PISA also identified a single salt bridge attributed to the light chain in proximity to residue G115, the first residue resolved in F1, placing the binding site near the furin cleavage site (**Table S1**). H8 binding induced a localized shift of the 396-405 loop, with a maximal displacement of 2.5 Å toward the Fab.

H8 also recognized the same topological site on postfusion F. Because domain II rotates relative to the rest of the protein during refolding around residue G433^27^, the Fab reoriented accordingly; whether H8 can remain bound through this transition is unclear. In the postfusion complex, contacts were again mediated mainly by the heavy chain, now comprising nine hydrogen bonds and two salt bridges, one per antibody chain (**Table S1**). The interface areas in pre- and postfusion were comparable (461 *vs.* 486 Å²). ΔG values for both conformations were estimated to be similar (−4.2 kcal/mol for prefusion and −3.1 kcal/mol for postfusion).

In summary, cryo-EM resolved Y10F as an apex-binding antibody that fits between glycan clusters on F2, confirmed the ability of Y10F to co-occupy F with 77.1 without major conformational changes of F, and showed that H8 targets a solvent-exposed loop in domain II that is accessible in both prefusion and postfusion conformations (see **Supplemental Fig.2** for antibody footprint on the F protein).

### In vivo efficacy and inhibition of fusion

We next evaluated the prophylactic efficacy of these mAbs *in vivo* using a cotton rat model of measles virus infection^29^ (**Fig. 5A**). Cotton rats were inoculated 14 hours before infection with a single dose (1 mg/kg) of the indicated mAbs. Four days after infection, lungs were collected and viral titers quantified (y-axis). A purified polyvalent human IgG (huIgG) preparation, which is used clinically for patients with primary immune deficiency and immune thrombocytopenia purpura, exhibited a neutralization titer of 640 at 60mg/mL and served as a control. As expected, mAb 77.1 confirmed its strong *in vivo* efficacy previously demonstrated^27^. The second most potent mAb was Y10F, which reduced viral titers close to the limit of detection. The other two mAbs (C6 and H8) performed comparably to huIgG, each reducing viral titers relative to PBS (diluent)-treated animals.

**Figure 5.**
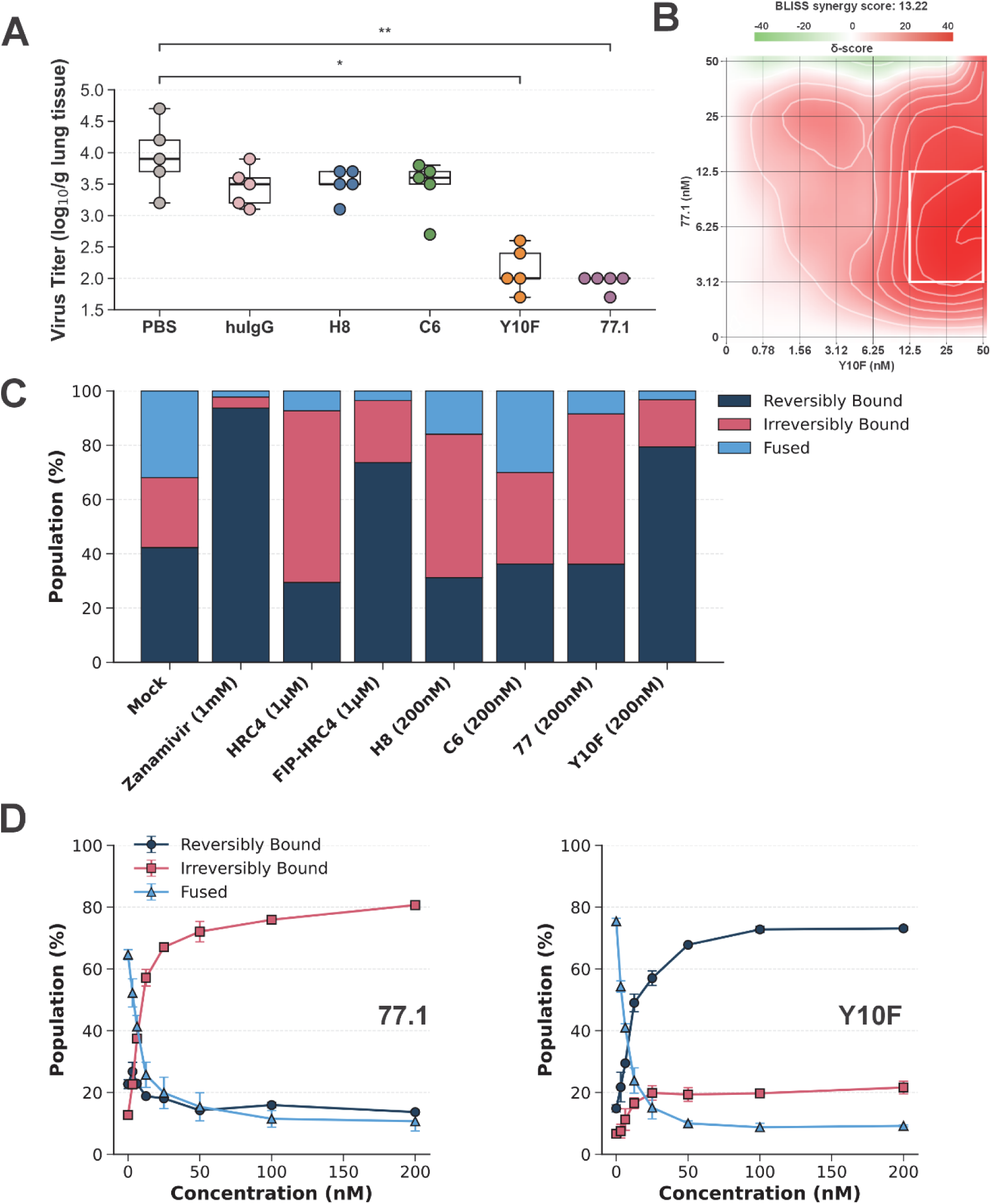
In Vivo Efficacy and Mechanism of Action of Anti-F Monoclonal Antibodies. **A)** Antibody transfer experiment to assess *in vivo* protective efficacy of mAbs. Cotton rats received intraperitoneal (IP) injection of 1 mg/kg the indicated mAbs or IgG purified from human sera (Carimune, CSL Behring) 14 hours before challenge with MeV. Animals in the control group received PBS before the challenge. Values correspond to MeV titers in lung tissue collected 4 days post-infection. Significance was shown using the Mann-Whitney-Wilcoxon test. P-value annotation legend: ns (non-significant): p <= 1.00, *: 0.01 < p <= 0.05, **: 0.001 < p <= 0.01, ***: 0.0001 < p <= 0.001. **B)** Synergistic inhibition of cell–cell fusion by 77.1 and Y10F antibodies. B3 H and F co-expressing effector cells were co-cultured with SLAM-expressing target cells in the presence of 77.1 and Y10F antibodies at the indicated concentrations (X- and Y-axes). Fusion was quantified after 18 hours using a β-galactosidase complementation assay, and synergy was assessed using the BLISS independence model. A moderate synergistic effect was observed, with a BLISS score of 13.22. The area of highest synergism is highlighted by a white rectangle. **C)** A bar chart summarizing the results of a mechanistic assay comparing various known inhibitors, including the newly described mAbs. For each condition, cells were categorized based on their interaction with red blood cells (RBCs): reversibly bound (dark blue) where the RBCs were only adhered by chimeric H and could be easily removed by treatment with zanamivir; irreversibly bound (pink) where RBCs were initially attached by chimeric H, but subsequent F protein refolding anchored the fusion peptide in the RBC membrane, making the RBCs attachment zanamivir-insensitive and retaining RBC on the HEK293T cell surface; and fused (light blue) where RBCs underwent membrane fusion with HEK293T cells expressing the fusion complex. The data are the results of three separate experiments (see **Supplemental** Fig. 5) **D)** Mechanistic assay over a range of mAb concentrations (x-axis), in which both 77.1 and Y10F efficiently block fusion (light blue triangles). 77.1 traps the F in the extended state (red squares). Y10F prevents F activation and F insertion (dark blue circles).

Since Y10F and 77.1 can simultaneously bind to the F protein (**Fig. 2B**), we tested whether they exhibit synergistic inhibition of fusion. Fusion was measured using an α/ω β-galactosidase complementation assay, where HEK293T target cells expressing SLAM and ω subunit were co-cultured with effector cells expressing MeV H/F and α subunit; and reconstituted enzyme activity was quantified by luminescence (**Fig. 5B**)^27^. Data were analyzed using SynergyFinder (https://synergyfinder.fimm.fi/) using the BLISS model. We observed detectable moderate synergism between Y10F and 77.1 in inhibiting viral fusion.

To define the stage of the fusion process at which each mAb acts, we used an assay measuring fusion of red blood cells (RBCs) with HEK293T cells expressing modified versions of MeV H and F (**Fig. 5C**)^30^. To distinguish F-mediated anchoring from H-mediated attachment, we employed a chimeric receptor-binding protein consisting of the MeV H stalk fused to the head domain of human parainfluenza virus type 3 hemagglutinin-neuraminidase (HPIV3 HN). This H-HN chimera allows releasable attachment to sialic acid receptors (via HN) while still permitting MeV H-mediated triggering of F. As expected, RBCs bound through HN-sialic acid interactions could be released by adding Zanamivir^27,30^. In untreated conditions, ∼38% of RBCs fused with HEK293T cells, ∼22% irreversibly bound via F (fusion initiated, but incomplete), and ∼40% were reversibly bound via HN only (fusion not initiated). Zanamivir treatment, which blocks HN-mediated attachment, prevented nearly all F-mediated anchoring, confirming that receptor engagement is required for F activation. Treatment with the HRC4^30^ peptide inhibitor yielded mostly irreversibly bound RBCs, indicating that F was activated, but fusion was blocked at a post-triggering step. By contrast, the FIP-HRC4^27,30^ peptide inhibitor prevented nearly all F-mediated anchoring, consistent with inhibition of F activation. MAbs 77.1 and H8 blocked fusion in the same manner as HRC4, with most RBCs irreversibly bound, confirming that these two mAbs permit (77.1) or promote (H8) the F extension and allow fusion peptide insertion into the target cell, but block completion of the F transition. The profile observed in the presence of C6 - at the concentration tested (200 nM)-was statistically indistinguishable from that of mock-treated samples, consistent with its lack of efficacy. In contrast, mAb Y10F behaves similarly to FIP-HRC4^30^, preventing the initiation of F activation. A dose–response version of this assay, with 77.1 and Y10F, confirmed that these antibodies act through distinct mechanisms: at higher concentrations, 77.1 drives cells into an irreversibly bound state with F trapped in an intermediate conformation, whereas Y10F drives cells into a reversibly bound state with F remaining untriggered (**Fig. 5D**).

### Viral evolution in the presence of the mAbs 77.1 and Y10F

We first evaluated the potential for viral escape from mAb 77.1; we subjected MeV to increasing concentrations of the antibody over five serial passages. Viral replication ceased after the fifth cycle, likely due to the accumulation of deleterious mutations. Nevertheless, viral populations from cycles four and five were harvested and subjected to deep sequencing to identify potential resistance-associated genetic changes. Remarkably, despite prolonged selective pressure from mAb 77.1, we detected no significant point mutations with allele frequency ≥10%, sequencing depth ≥20 (data not shown).

We next repeated the viral evolution experiment with both 77.1 and Y10F (since these two antibodies were the most potent *in vivo*, as shown in **Fig. 5A**), using a different strategy. Instead of progressively increasing antibody concentrations, viruses were passaged for over 50 days in alternating concentrations of mAbs to permit ongoing replication (see Materials and Methods). This approach yielded eight distinct viral populations: four selected in the presence of 77.1 and four in the presence of Y10F. Three of the four populations grown with 77.1 showed no mutations in the F protein (**Supplemental Fig. 3**), while one population contained a single amino acid substitution (M46V) (**Fig. 6A**). In contrast, all four populations selected with Y10F acquired one or more amino acid substitutions in F (**Fig. 6A**). Among these, only the F375S variant reached complete fixation (100% allele frequency), whereas the others displayed partial penetrance. Nevertheless, all Y10F-selected populations were resistant to Y10F inhibitory activity but retained sensitivity to 77.1, albeit with reduced potency compared to the parental virus (**Supplemental Fig. 4**). The M46V (selected in the presence of 77.1) was also resistant to Y10F and still inhibited by 77.1. None of these mutations mapped directly to the binding interface defined in **Figs. 2** and **3**, suggesting that viral evolution under antibody pressure may favor allosteric changes or modifications of F protein function that diminish antibody efficacy (**Fig. 5B**).

**Figure 6.**
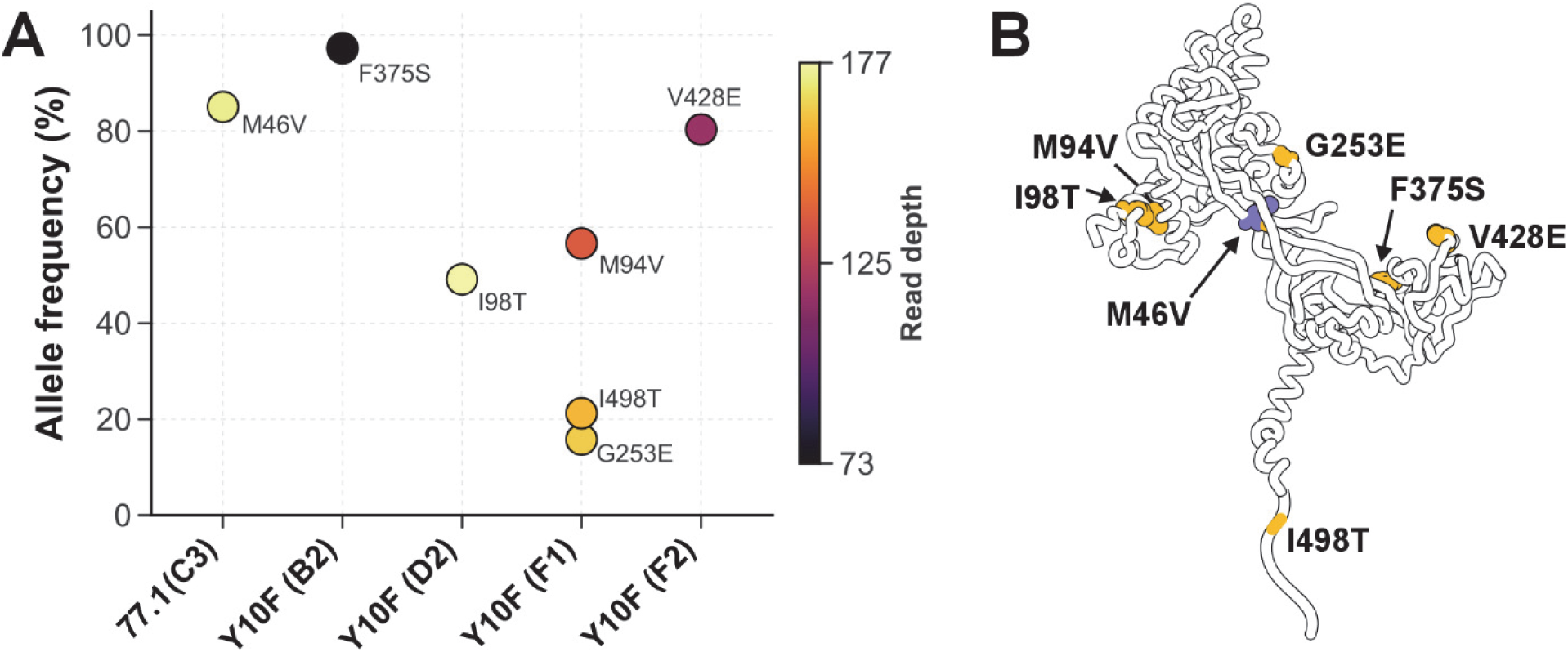
Viral Evolution Under Selective Pressure of 77.1 and Y10F mAbs. **A)** Viral evolution assay results from control (mock), as well as passages in the presence of Y10F and 77.1, with four replicates per mAb, were evaluated for mutations in the F gene. Only mutations with an allele frequency greater than 10% and a minimum read depth of 20 were included in the analysis. **B)** Location of point mutations identified in A) mapped onto a single protomer of the prefusion F structure. Mutated residues are highlighted, including I498T, which is located in the transmembrane region and shown schematically.

## Discussion

The measles virus (MeV) vaccine is highly effective but cannot be given to immunocompromised or pregnant individuals, leaving them dependent on herd immunity^1,31,32^. This protection is increasingly fragile due to suboptimal coverage and MeV’s extreme transmissibility (R₀ = 12–18), driving resurgences in the U.S. and worldwide. MeV still causes 10–20 million infections and >200,000 deaths annually, compounded by immune amnesia that erodes immunity to other pathogens^1–6,33–36^. No licensed antivirals exist, and current post-exposure prophylaxis with immune sera shows declining potency^7–15^. While vaccine-induced antibodies primarily target hemagglutinin, natural infection elicits potent antibodies against the fusion protein, highlighting F as a promising therapeutic target^19,20,37–39^.

In this study, we investigated the neutralizing activity of mAbs against the MeV F protein, generated in mice and humanized by grafting their variable regions (Fab) onto human constant domains (Fc). One mAb, 77.1, has been previously described both functionally and structurally^27^; it binds an epitope present in both the prefusion and extended conformations of the F protein, and functions by arresting the F conformational transition at that intermediate step. Here, we evaluated its potential for use in combination therapies and its barrier to viral resistance. A second mAb, Y10F, has previously been shown to possess neutralizing activity^40,41^, but its sequence, epitope specificity, and mechanism of action were unknown. The other two mAbs, H8 and C6, were selected for their distinct binding properties: H8 recognizes both pre- and post-triggered forms of F, whereas C6 binds only a post-triggered epitope.

The three neutralizing mAbs (Y10F, 77.1, and H8) demonstrate that the MeV F protein contains multiple vulnerable epitopes that can be effectively targeted. Y10F binds an apical prefusion epitope, prevents F activation by stabilizing its prefusion state, and neutralizes efficiently both *in vitro* and *in vivo*. However, viral escape can occur through allosteric mutations outside the binding interface. 77.1 recognizes F in both pre- and post-triggered states, blocks fusion at a late stage, and exhibits a high barrier to escape. H8 binds epitopes present in both conformations and, uniquely, promotes F activation upon binding a previously undescribed mechanism of action. This binding profile is similar to that of the 1H1 antibody, which targets the Nipah virus F protein at an analogous epitope on the outer side of domain II^42^. Like H8, 1H1 is also a neutralizing antibody and recognizes both the prefusion and postfusion states of the F protein. However, 1H1 functions through a conventional blocking mechanism and does not trigger F protein activation. This confirms that the premature activation induced by H8 represents a unique mechanism among this class of antibodies. In addition, H8 neutralizes a broad spectrum of viruses, including hyperfusogenic variants from neuropathogenic isolates^28^ and blocks fusion mediated by all the F variants obtained from the viral evolution with Y10F and 77.1 (data not shown). By contrast, C6, which recognizes only a post-triggered epitope in F, failed to block viral entry or fusion, underscoring the importance of antibodies that target pre- and intermediate conformations.

Collectively, these findings establish the MeV F glycoprotein as a rich and underexplored source of neutralizing epitopes. A cocktail of long-acting, humanized anti-F antibodies, such as Y10F and 77.1, could provide durable protection with a high barrier to resistance, complement vaccination, and address a critical therapeutic gap for individuals who are unvaccinated, vaccine-ineligible, or have waning immunity^1–6,19,20,37–39^.

## Acknowledgements

The authors thank NIAID R56AI183536, NIAID R01AI176833 and U19AI181984 (MP), NINDS R01NS105699 and R01NS091263 (MP), NIAID R21AI180456 (EOS), Swiss National Science Foundation Postdoc Mobility fellowships P2EZP3_195680 (DSZ) and P500PB_210992 (DSZ), and Institutional Funds of the La Jolla Institute for Immunology (EOS) for funding, Measles biobank (https://ciri.ens-lyon.fr/teams/IbIV/measles-biobank) for funding, and Dr. Ruben Diaz and the LJI Cryo-EM Core for data collection. Instrumentation in the cryo-EM core was supported by U19 AI109762-S1, gifts from the GHR Foundation and private philanthropic support. Molecular graphics and analyses performed with UCSF ChimeraX, developed by the Resource for Biocomputing, Visualization, and Informatics at the University of California, San Francisco, with support from National Institutes of Health R01-GM129325 and the Office of Cyber Infrastructure and Computational Biology, National Institute of Allergy and Infectious Diseases. We thank N. Pascual and C. Leedale for technical assistance. We thank Dr. Rik de Swart for the critical reading of the manuscript.

## Material and Methods

### Plasmids

The genes of MeV B3, D8, Vaccine hemagglutinin, fusion protein, and CD150 (or hSLAM) proteins were sub-cloned into the mammalian expression vector pCAGGS. The identification codes of the different fusion proteins are: B3-F (GS63325-1), D8-F (GS77420-3), Moraten-F (GS55588-1), L454W-F (GS55608), and E455G-F (GS74446-1). A codon-optimized fusion protein sequence derived from the Measles virus strain Ichinose-B95a (Taxonomy ID: 645098) was synthesized by GeneScript and subcloned into the pcDNA3.1 vector. Stabilizing point mutations (E170G and E455G) previously characterized in SSPE and MIBE Measles strains^26,28,43^ were sequentially introduced using the Q5 Site-Directed Mutagenesis kit. The ectodomain construct was generated by removing the transmembrane region and cytoplasmic tail residues (amino acids 496-550) using the Q5 Site-Directed Mutagenesis kit. To generate Drosophila S2 stable cell lines, fusion protein ectodomain construct E170G/E455G was subcloned from the pcDNA vector to the pMT-Puro vector, including a C-terminal enterokinase cleavage site-twin-strep-tag via NEB HiFi DNA assembly.

### Antibodies

MeV F–specific murine monoclonal antibodies 77.4 and Y10F were generated as described previously^41^. Chimeric antibody 77.1 has been functionally and structurally characterized^27^. The remaining two antibodies, H8 and C6, were derived from murine hybridomas obtained after immunization with stabilized MeV F; their Fab regions were subsequently sequenced and cloned into a human Fc backbone, as described for 77.1^27^ ***Cells.*** HEK293T (human kidney epithelial), Vero, and Vero-hSLAM/CD150 (African green monkey kidney) cells were grown in Dulbecco’s modified Eagle’s medium (DMEM; Life Technologies; Thermo Fisher Scientific) supplemented with 10% fetal bovine serum (FBS, Life Technologies; Thermo Fisher Scientific) and antibiotics at 37°C in 5% CO_2_. The Vero-hSLAM/CD150 culture media were supplemented with 1mg/mL Geneticin (Thermo Fisher Scientific). Drosophila S2 cells were cultured in complete Schneider’s Drosophila Medium (Gibco) supplemented with 10% heat-inactivated fetal bovine serum (FBS) and 1% penicillin-streptomycin (100 U/mL penicillin and 100 µg/mL streptomycin) at 25°C without CO_2_ or Lonza Insect Xpress Medium. ExpiCHO cells were maintained in ExpiCHO Expression Medium (Gibco) supplemented with 8 mM L-glutamine in a humidified incubator at 37°C with 8% CO_2_.

### Viral rescue

HEK293T cells were transfected using Lipofectamine 2000 on 6-well plates coated with poly-D-lysine. The transfection mixture included purified plasmid constructs coding for viral proteins N (GS58900-1), P (GS58943), and L (GS58944), all sourced from Epoch Life Science according to our design specifications. Additionally, purified plasmids for T7 polymerase (GS58929) and the genome for the MeV B3 mCherry wt virus bearing either the B3 H and F (GS69677-10), or the D8 H and F (GS77420-2), or the H and F from the Moraten vaccine strain (GS77420-1), or the B3 H and F-L454W (GS69677-1) were included. Cells were incubated overnight at 37°C in Opti-MEM medium. The medium was then replaced with warm DMEM supplemented with 10% FBS. After 1 hour, cells were subjected to a heat shock treatment by incubating them at 42°C in a water bath for 3 hours. Subsequently, the cells were returned to a 37°C incubator for 48 hours. After this incubation, the HEK293T cells were detached and transferred to Vero-hSLAM/CD150 cells in a 6-well plate and incubated at 32°C for 5 days. RNA from the virus sample was then sequenced, and the virus stock was used for subsequent in vitro and in vivo experiments (see^27^).

### In vitro viral entry

Vero-hSLAM/CD150 cells were plated in 96-well plates (1 × 10^4^ cells/well). The following day, cells were treated with serial dilutions of the indicated antibodies. Cells were then infected with the indicated viruses for 3 h at 37°C (200-300 pfu/well). After 3h, the medium was replaced with medium containing 2% of methylcellulose 2× complete medium (1:1). After 24 h, fluorescent/infected cells were counted using a Cytation 5 (BioTek) and analyzed with Gen5 3.11 software (min object size 25 µm, max object size 135 µm, threshold 3,000). The samples were tested in three separate experiments performed with technical duplicates in five-fold dilutions in 96-well plates. Data obtained from each plate was analyzed in OriginPro 2023, where all biological and technical replicates were fitted with a logistic regression function with shared IC_50_ and Hill slope values, and A1 and A0 (highest and lowest signals) were used to normalize the data. Based on the normalized data, the percent of inhibition was calculated using the following equation: [1 - X]*100, where X is the fraction of the normalized signal.

### Beta-Galactosidase (Beta-Gal) complementation-based fusion assay

The Beta-Gal complementation-based fusion assay was performed as previously described^30^. In brief, HEK293T cells transiently transfected with the hSLAM/CD150 receptor and expressing the ω subunit of β-galactosidase were co-cultured with cells co-expressing the MeV glycoproteins F and H together with the α subunit of β-galactosidase. Cell fusion results in α–ω complementation, which was measured in the presence or absence of inhibitory compounds. After 18 h, cells were lysed to terminate the fusion process, and fusion levels were quantified using the Tropix Galacto-Star™ chemiluminescent reporter assay system (Applied Biosystems) and read on a Tecan M1000 microplate reader. Biological replicates were recorded and globally fitted with a logistic function in OriginPro 2023. The obtained parameters were used to normalize the data per sample and plotted with standard error calculated from all data points.

### Cell surface staining with conformation-specific anti-F mAbs

HEK293T cells transiently transfected with viral glycoprotein constructs were incubated overnight at 37°C in complete medium (DMEM supplemented with 10% FBS). The following day, cells were shifted to the indicated temperatures and incubation times (as shown in the figures). To detect specific MeV F conformations, cells were incubated on ice for 1 h with chimeric monoclonal antibodies (mAbs), followed by a 1 h incubation on ice with Alexa Fluor 594 or Alexa Fluor 488-conjugated anti-human secondary antibody (1:1000; Life Technologies). Cells were then fixed on ice with 4% paraformaldehyde for 10 min and counterstained with DAPI (1:1000; Thermo Fisher). Imaging was performed using a Cytation 5 system, and relative fluorescence of antibody-bound cells was quantified with BioTek Gen5 software.

### Viral evolution assay

Selection of Measles Virus Escape Mutants in the Presence of Monoclonal Antibodies Cell Culture and Virus Infection.

### Phase 1: Cell Culture and Virus Infection

Vero-hSLAM/CD150 cells were seeded in two 96-well plates and infected with 100 plaque-forming units (PFUs) per well of recombinant Measles virus (MeV) B3 wild-type expressing mCherry. Infections were carried out in Opti-MEM (Gibco, 31985070) supplemented with 10% penicillin-streptomycin. After 2 hours, the infection medium was replaced with serially diluted concentrations of monoclonal antibodies Y10F and 77.1, starting at 10 μg/mL and diluted 1:2 down to 0.04 μg/mL.

### Phase 2: Passaging Under Antibody Pressure

On day 10 post-infection, wells showing evidence of ongoing viral replication in the presence of highest (*i.e.,* 10 μg/mL) antibody concentration (1-77.1: B1 and G1; 1-Y10F: F1 and E1) were selected. The contents of each well were transferred to T75 flasks containing Vero-hSLAM/CD150 (VS) cells in Opti-MEM supplemented with 10% penicillin-streptomycin and 10 μg/mL of the corresponding antibody.

Due to limited infection after 3 days, additional wells from the antibody selection plates were chosen and expanded: wells 2-77.1 (A3, C3, E3) and 2-Y10F (B2, D2, F2) were transferred to T75 flasks and cultured in Opti-MEM with 10% pen-strep, containing reduced antibody concentrations (1.25 μg/mL for 77.1; 2.5 μg/mL for Y10F).

On day 15, the medium from the initial T75 flasks (*i.e.,* 1-77.1: B1 and G1; 1-Y10F: F1 and E1) was replaced with Opti-MEM + 10% pen-strep without antibody. Virus supernatants were collected and frozen at −80°C on day 16. Antibody concentrations in the second batch flasks were further reduced on day 17 due to a lack of visible infection (0.625 μg/mL for 77.1; 1.25 μg/mL for Y10F). Viral supernatants from these cultures were collected on days 23 and 28 (2-Y10F: B2, D2, F2; 2-77.1: A3, C3, E3), frozen, and subsequently used for further passages.

### Sequential Antibody Reduction and Virus Harvesting

*Phase 3*: On day 28, all remaining samples were subsequently used to infect T75 flasks of VS cells at 10-fold lower antibody concentrations (0.125 μg/mL for 2-Y10F, 0.0625 μg/mL for 2-77.1, and 0.5 μg/mL for first-round isolates). Viruses were all harvested in the following 10-12 days.

*Phase 4*: On day 39, these eight virus stocks were then used to infect new T75 flasks of VS cells. After three hours, antibody was reintroduced at 2-times lower concentrations (0.0625 μg/mL for 2-Y10F, 0.0313 μg/mL for 2-77.1, and 0.25 μg/mL for 1-Y10F and 1-77.1).

*Phase 5*: On day 54, each virus was collected, titered, and sequenced.

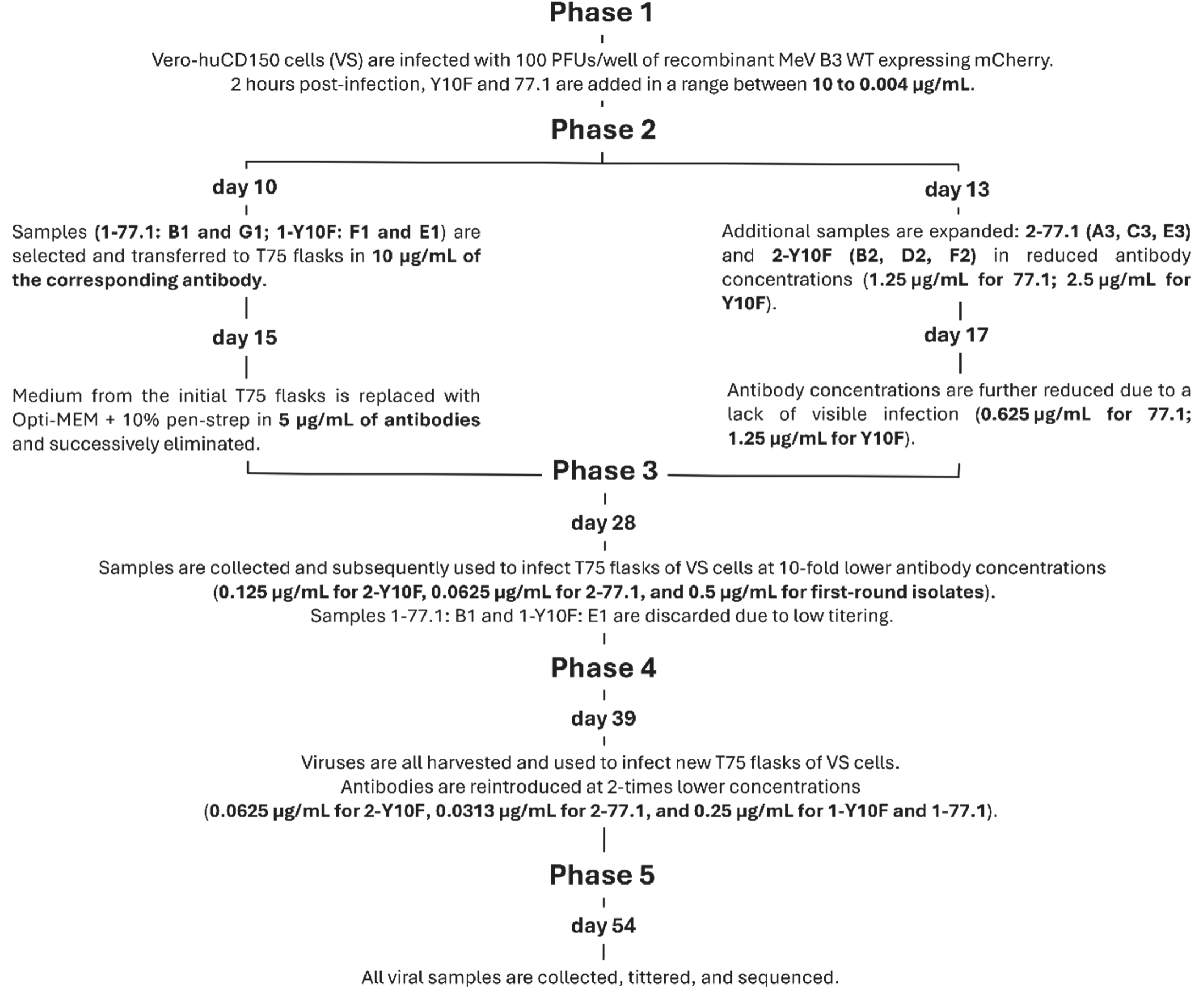

### In vivo experiments: Cotton rats

Inbred cotton rats (*Sigmodon hispidus*) were purchased from Envigo, Inc., Indianapolis. Both male and female cotton rats aged 5 to 7 weeks were used. For i.n. infection, 10^5^ TCID_50_ of MeV (strain B3 wt EGFP) was inoculated intranasally to isoflurane-anesthetized cotton rats in a volume of 100 µL. Animals received either the indicated amount of the indicated antibodies, purified human IgG (Carimune, CSL Behring, measles neutralizing titer of 640 at 60 mg/mL), or vehicle 14 hours prior to infection. Four days after infection, the animals were euthanized by CO_2_ inhalation, and their lungs were collected and weighed. Lung tissue was homogenized with a glass Dounce homogenizer. Serial 10-fold dilutions of supernatant fluids were assessed on Vero-hSLAM/CD150 cells for the presence of infectious virus in 48-well plates using cytopathic effect (CPE) as the endpoint. Plates were scored for CPE microscopically after 7 days, and the tissue culture infectious dose 50 (TCID_50_) was calculated. The results are presented as TCID_50_/g of lung tissue.

### RBC fusion assay

RBC fusion assays were performed using HEK293T cells transiently expressing the MeV H_Y17H–HPIV3_T193A chimera and MeV F (T461I). Cell monolayers were washed three times with serum-free medium and incubated at 4°C for 30 min in CO₂-independent medium containing 1% RBCs. Cells were then exposed to varying concentrations of inhibitors and shifted to 37°C for 60 min.

To quantify RBC binding and fusion, zanamivir was added at a final concentration of 10 mM, and the cells were incubated at 4°C for 30 min with gentle agitation. The supernatant was collected into V-bottom 96-well plates to measure **reversibly bound RBCs**. The cells were subsequently incubated at 4°C for 30 min in ACK lysing buffer, and the supernatant was collected into V-bottom 96-well plates to quantify **irreversibly bound RBCs**. Finally, the remaining cells were lysed in lysis buffer and transferred to flat-bottom 96-well plates to determine **fused RBCs**. Absorbance at 410 nm was measured using a Tecan My1000PRO microplate reader to quantify RBCs in each category. Assays were performed in CO_2_-independent media at pH 7.2 for the bar graphs in **Fig. 5C** and **Supplemental Fig. 5,** and at pH 7.6 for the dose response curves in **Fig. 5D**.

### F_ECTO_ purification

Fecto was purified as previously described from Drosophila S2 cells^27^. Protein concentration was estimated using the molar extinction coefficient at 280 nm (48,820 M⁻¹ cm⁻¹).

### Antibody expression

Monoclonal antibodies were produced in ExpiCHO cells using the manufacturer’s high-titer transfection protocol with pTRIOZ-mAb plasmids encoding the antibody heavy and light chains. After seven days, cells were removed by centrifugation at 4,000 × g, and the clarified medium was adjusted to pH 8.0 with 1 M HEPES (pH 8.0). Antibodies were captured by incubating the medium overnight at 4°C with protein A agarose. The next day, beads were washed with 20 mL HBS, and bound antibody was eluted with 15 mL 0.2 M glycine (pH 2.0). The eluate was immediately neutralized by adding 10% (v/v) 1 M HEPES (pH 8.0).

For the Fab generation, purified IgG at 1 mg/mL was digested with 2% (w/v) cysteine-activated papain for 4 h at 37°C in a water bath. The reaction was quenched by adding 0.5 M iodoacetamide to a final concentration of 50 mM. The mixture was diluted tenfold with 10 mM HEPES (pH 8.0) to reduce ionic strength and loaded onto a 1 mL Mono S column pre-equilibrated with 20 mM HEPES (pH 8.0). Fab fragments were collected in the flow-through, whereas Fc fragments were eluted in a single step with 20 mM HEPES (pH 8.0) containing 2 M NaCl. Isolated Fabs were concentrated, buffer-exchanged into HBS, and stored at 4°C. Protein concentrations were determined from A280 using the calculated molar extinction coefficients.

### Surface plasmon resonance

Binding experiments were performed on a Carterra LSA instrument. Purified monoclonal antibodies (mAbs) were covalently immobilized onto an HC30M sensor chip at a concentration of 2.5 µg mL⁻¹, with each antibody injected in duplicate. A serial dilution of the analyte (F_ECTO_) was applied to the chip in ascending order: 3.125 nM, 6.25 nM, 12.5 nM, 25 nM, 50 nM, and 100 nM. The resulting binding responses were recorded over time. Kinetic parameters were extracted by globally fitting the sensorgrams to Equations 1–6 using OriginPro software.

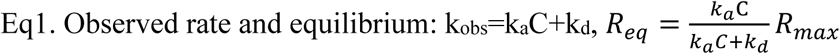

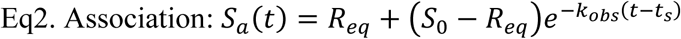

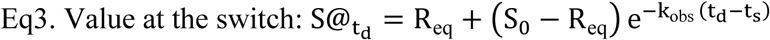

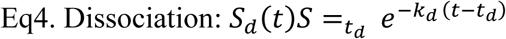

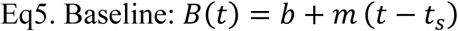

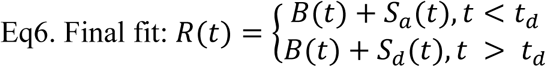

Where: t: time (s). t_s_: association start (s). t_d_: dissociation start (s), C: analyte concentration (M), k_a_: association rate constant (M^−1^ s^−1^), k_d_: dissociation rate constant (s_−1_), k_obs_: observed association rate (s^−1^), R_max_: maximum response at full occupancy (response units), R_eq_: equilibrium response at C (response units), S_0_: residual bound signal at t_s_ (response units), S_@td_: signal at the switch to buffer (response units), b: baseline offset (response units), m: linear drift (response units s^−1^)

### Grid preparation

#### Negative stain

Negative-stain electron microscopy was performed using 400-mesh copper grids coated with a continuous carbon film (CF400-Cu; Electron Microscopy Sciences). Prior to sample application, the grids were rendered hydrophilic by glow-discharging them in an easyGlow Glow Discharge Cleaning System (Pelco) for 15 s at 15 mA. Protein samples were diluted to an absorbance of 0.01 at 280 nm; 3 µL of this solution was then deposited onto the grid and incubated for 1 minute. The excess sample was removed by blotting, and the grid was washed three times with 3 μL of deionized water, with blotting performed between each wash. Negative staining was performed by applying 3 µL of a 1 % uranyl acetate solution to the grid, blotting twice, then adding an additional drop of stain and incubating for one minute before a final blot to remove excess stain. The grids were allowed to air-dry prior to imaging.

#### Cryo-EM sample preparation and data acquisition

Complex stoichiometries were guided by NS-EM. Except where noted, complexes were incubated with a twofold molar excess of Fab relative to the number of F binding sites. Accordingly, samples were prepared at trimer:Fab ratios of 1:2 for F_ECTO_:Y10F (one F trimer to two Y10F Fabs) and 1:2:6 for F_ECTO_:Y10F:77.1 (twofold excess of 77.1). For H8, Fab was added at a 1:1 ratio per F binding site (1:3 trimer:H8), because this complex was not purified before imaging.

F_ECTO_:Y10F and F_ECTO_:Y10F:77.1 complexes were purified by size-exclusion chromatography (SEC) before grid preparation. The F:H8 complex was applied directly to grids after a 15-minute incubation to minimize postfusion conformational transition that could occur during SEC. Data were collected on graphene oxide (GO) grids as described previously^27^. All F_ECTO_-Fab complexes were recorded on Quantifoil R2/1 or R2/2 grids covered with GO grids prepared as described previously^27^. Protein with 0.25-35 absorbance values at 280 nm was placed on grids and blotted for 2-3s. Grids were loaded to the FEI Titan Krios G3 autoloader and screened for ice quality, graphene oxide layer quality, and particle distribution. Movies were recorded with an exposure rate of 70 e/Å², with 50 frames per movie saved as non-gain normalized LWZ TIFF files. Collection statistics per dataset are reported in **Table S2**.

#### Cryo-EM data processing

All datasets were processed in cryoSPARC v4.5 or v4.6 using a standard workflow with dataset-specific adjustments during late refinement. Movies were corrected for beam-induced motion with Patch Motion, and defocus values were estimated with Patch CTF in cryoSPARC Live. Particles were initially picked with the blob picker using an intentionally broad particle diameter (80–120 Å) to over-pick F_ECTO_–Fab complexes. Particles were first extracted and Fourier-cropped to 64 pixels to speed up early processing.

Particle stacks were extensively cleaned by 2D classification following the Deep2D protocol described previously^27^, and class averages showing well-defined protein features were retained. Two independent ab initio reconstructions were run per dataset, one imposing C1 and one C3 symmetry, each with five classes; for the Y10F dataset, only C1 was used. The best ab initio models were then used for heterogeneous refinement against all classes, iterating until discrete 3D classes emerged. The best-looking class or classes were pooled and subjected to non-uniform refinement (NUR), after which particles were re-extracted with larger Fourier-cropped boxes (160–256 pixels) for further NUR and additional 3D classification to improve compositional and conformational homogeneity.

For final map generation, only particles contributing to the best-resolved class were retained and re-extracted with a Fourier crop chosen so that the 0.143 FSC resolution fell within approximately three-quarters of the box frequency (to avoid box-size–limited resolution). Final refinements included NUR, symmetry expansion where appropriate, focused/local refinement with masks, and targeted 3D classification, followed by a final round of local refinement. The final particle set was subjected to reference-based motion correction and both global and local CTF refinement, after which a final local refinement produced the maps reported.

#### Model building

Model building proceeded by rigid-body fitting of the previously reported F_ECTO_ structure (^27^PDB 8UUP) into each cryo-EM map. Antibody Fab fragments were modeled with AlphaFold2 and docked into the density using UCSF ChimeraX^44,45^. The combined F_ECTO_–antibody models were then iteratively adjusted manually in Coot^46^, followed by cycles of real-space refinement in PHENIX^47^ and additional manual corrections in Coot.

#### Interface interactions determination

Interface contacts, buried surface area, and estimated binding energy were calculated with PISA (CCP4 suite; ^48^). Interaction counts are reported as unique contacts from all symmetry-related interactions.

#### Formation of F_ECTO_-Fab Complexes for Determining Hydrodynamic Radius and Differential Scanning Fluorimetry

F_ECTO_ (F_ECTO_ E170G, E455G, residues 1-495) was mixed to a final concentration of ∼1.7 µM (trimer) and incubated with Fab at room temperature for 15–30 min. Fab was added at a threefold molar excess for 77.1, H8, and C6, and at a onefold molar excess for Y10F (ratios calculated relative to the F_ECTO_ trimer). The resulting mixtures were used directly for downstream analyses.

#### nanoDSF and DLS measurements

Thermal unfolding of F_ECTO_ with or without Fab, and of Fab alone, was measured on a NanoTemper Prometheus Panta. F_ECTO_–Fab complexes were assembled 30 min before acquisition and incubated at room temperature. For each experiment, three capillaries contained the F_ECTO_–Fab complex, two capillaries contained the corresponding Fab alone, and one capillary contained HBS as a buffer control. Data were collected with a 1°C/min temperature ramp and 100% excitation laser power. Unfolding profiles are reported as the first derivative of the intrinsic fluorescence at 330 nm. Dynamic light scattering was recorded in parallel, and the cumulant hydrodynamic radius was calculated and plotted as a function of temperature.

#### Next Generation Sequencing and variant calling

Viral whole-genome sequencing of MeV samples was performed using a metagenomic next-generation sequencing approach and a previously described protocol^49^ Libraries were sequenced on NextSeq 2000 with 2x150bp or 1x100bp read format. Raw reads were trimmed and quality filtered with fastp (v0.23.4)^49^ with the parameters --cut_mean_quality 20 --cut_front --cut_tail -- length_required 20 --low_complexity_filter --trim_poly_g --trim_poly_x. For variant calling, filtered reads were processed with the RAVA workflow (default parameters) using the MeV plasmid map as a reference (https://github.com/greninger-lab/RAVA_Pipeline/tree/2025-08-18_DZ_MP)^50^, (https://doi.org/10.1101/2019.12.17.879320). Raw sequencing data are publicly available in NCBI BioProject PRJNA1307578.

#### Statistical analysis

Data analysis and visualization were performed in Python using NumPy, pandas, SciPy, Matplotlib, seaborn, and statannotations. Unless stated otherwise, group comparisons used two-sided Mann– Whitney–Wilcoxon tests. When multiple pairwise comparisons were made within a figure panel, p-values were adjusted with the Holm–Bonferroni method to control the family-wise error rate. Exact p-values and test statistics (U) are reported in the figure legends. P-value annotations on plots follow this legend: ns, 5.00×10⁻² < p ≤ 1.00×10⁰; *, 1.00×10⁻² < p ≤ 5.00×10⁻²; **, 1.00×10⁻³ < p ≤ 1.00×10⁻²; ***, 1.00×10⁻⁴ < p ≤ 1.00×10⁻³; ****, p ≤ 1.00×10⁻⁴.

#### Chemicals and peptides

*N*-(3-cyanophenyl)-2-phenylacetamide (also known as 3G) was commercially acquired from ZereneX Molecular Limited (UK). The purity of 3G was tested by high-pressure liquid chromatography (HPLC) and shown to be >95% pure.

Peptides were purchased from CPC, Ltd. Bromoacetyl cholesterol and bismaleimide cholesterol derivatives were custom made by Charnwood Molecular Ltd. Dimethylsulfoxide (DMSO), tetrahydrofuran (THF), *N,N*-diisopropylethylamine (DIPEA) were purchased from Sigma Aldrich, bismaleimide-PEG_11_ from Quanta BioDesign. HPLC purification was performed on a 1100 Series Agilent HPLC system equipped with a UV diode array detector and a fraction collector using a reverse phase (RP.) Phenomenex Jupiter C4 LC column 300Å (150 x 21.2 mm, particle size 5 μm). MALDI-TOF analysis was performed on a Bruker UltrafleXtreme MALDI-TOF instrument.

## Data availability

Cryo-electron microscopy density maps and atomic coordinates have been deposited in public repositories. The Y10F–F ectodomain complex is available in the PDB (9Q5M) and EMDB (EMD-72240); the 77.1–Y10F–F ectodomain complex is available in the PDB (9Q5V) and EMDB (EMD-72242); the H8–F ectodomain complex in the prefusion state is available in the PDB (9Q7B) and EMDB (EMD-72293); and the H8–F ectodomain complex in the postfusion state is available in the PDB (9Q71) and EMDB (EMD-72283).

## Contributions

DSZ: Cloned and expressed proteins; recorded all negative stain and cryo-EM data; processed and solved structures; wrote the code for analysis; analyzed all data presented in the manuscript; prepared figures; wrote the manuscript. RDM: Performed virological and cell biology experimentation; monoclonal antibodies development; analyzed data; prepared figures; contributed to manuscript writing. DL: Performed cell-based and virological experiments; analyzed the data; prepared figures.GN: Assisted with screening, protein purification, and negative stain electron microscopy (NS-EM). GZ: Performed virological and cell biology experimentation; analyzed data. LDC: Rescued recombinant viruses; performed virological experiments; analyzed data. GJT: monoclonal antibodies development. GK: hybridoma production and monoclonal development. MA: Performed monoclonal antibodies binding analysis. GL: Performed virological studies; analyzed data. DV: sequencing of viral population; data analysis. KMH: Analyzed data; contributed to manuscript writing and figure preparation. BH: Monoclonal antibodies development; analyzed data. ALG: data analysis and manuscript writing. SN: Conducted in vivo experimentation, data analysis, and interpretation; supervised the work; contributed to manuscript writing. EOS: Analyzed and interpreted results; supervised the work; secured funding; contributed to manuscript writing. MP: Designed experiments; monoclonal antibodies development; performed virological and cell biological experimentation; analyzed and interpreted results; supervised the work; secured funding; wrote the manuscript.

## Ethics declarations

BH, MP, DSZ, and EOS have a provisional patent related to the antibodies as passive immunotherapy. All the authors declare that they have no competing interests in relation to the work described in this manuscript.

---

**Figure S1.**
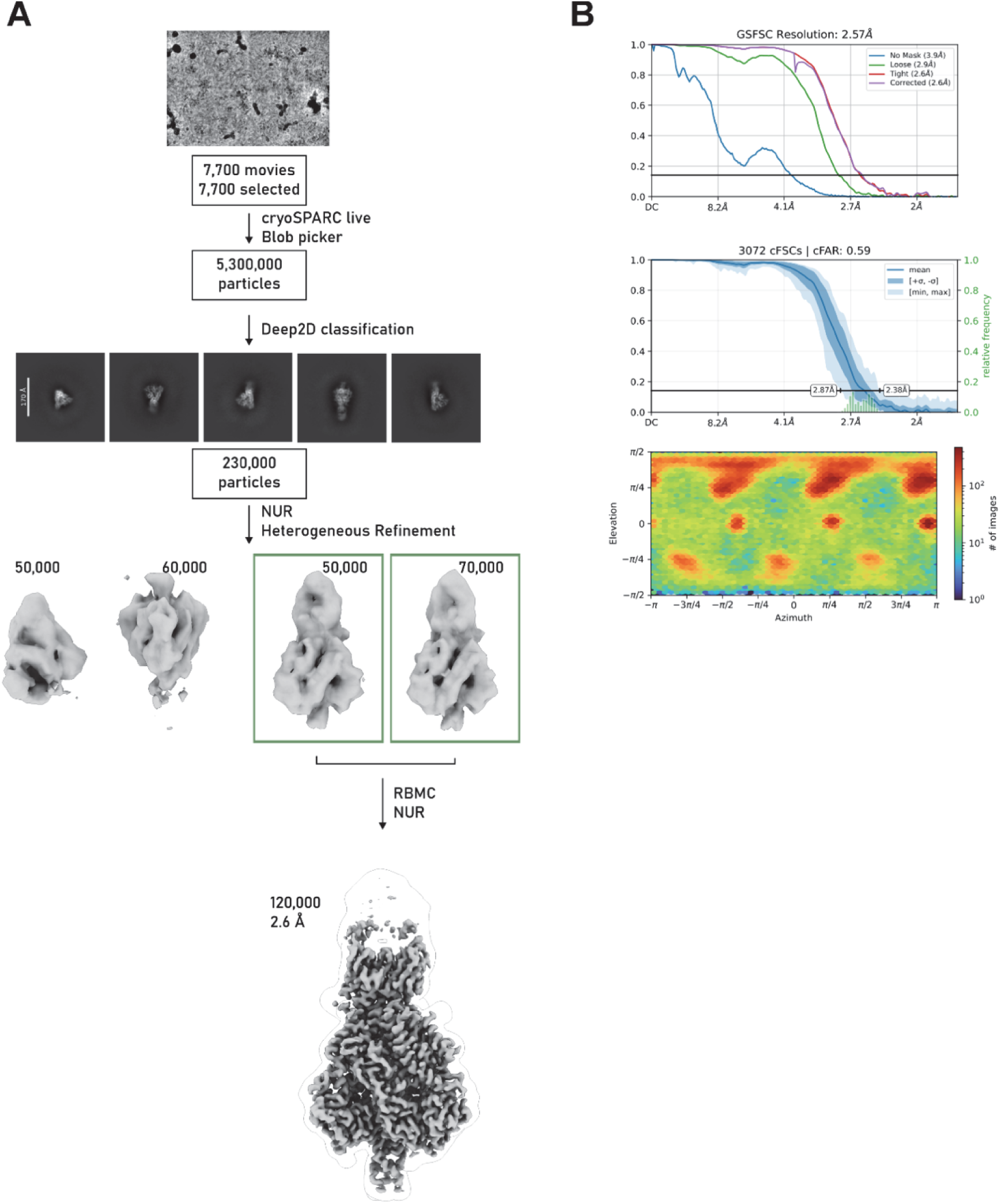
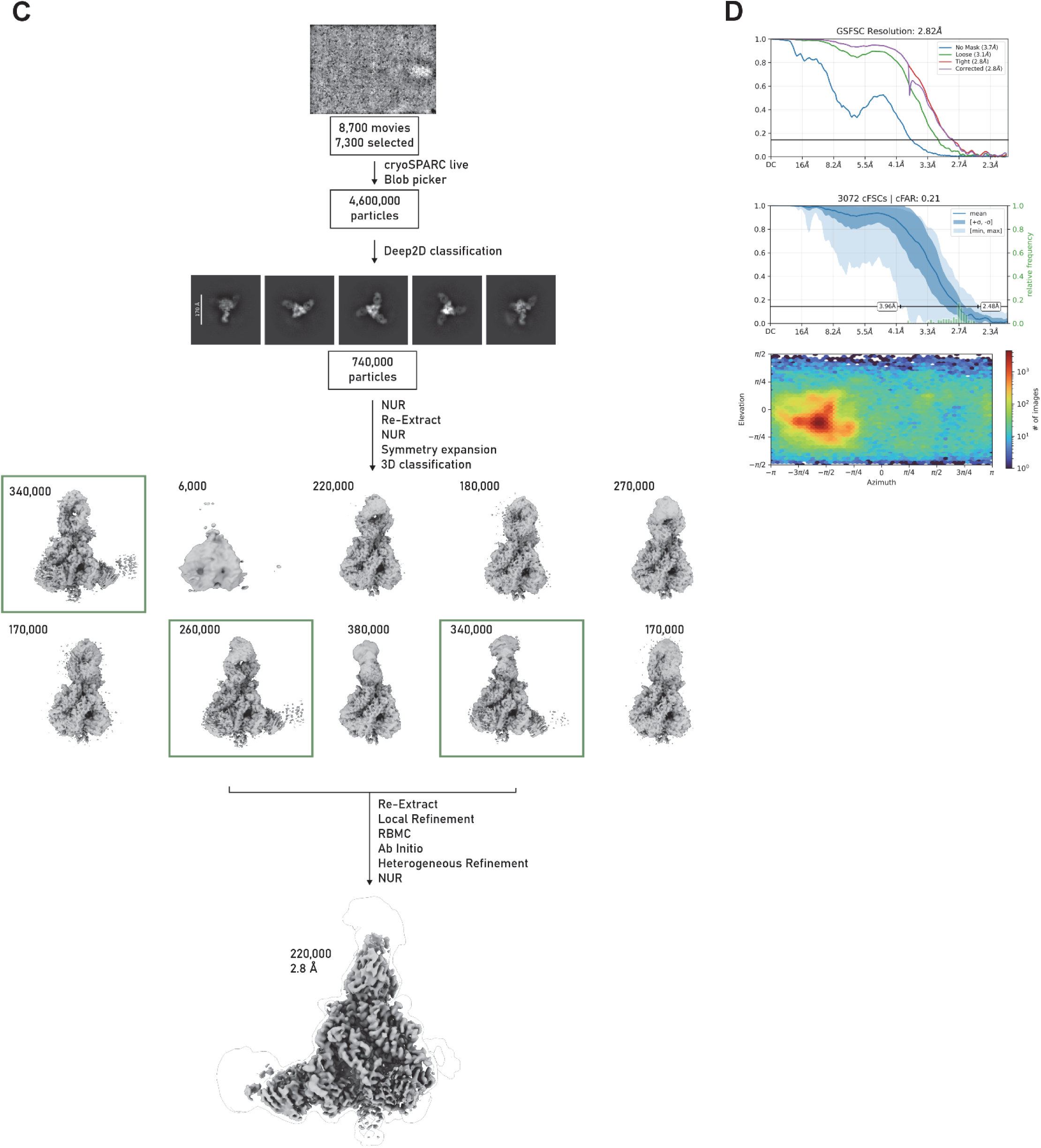
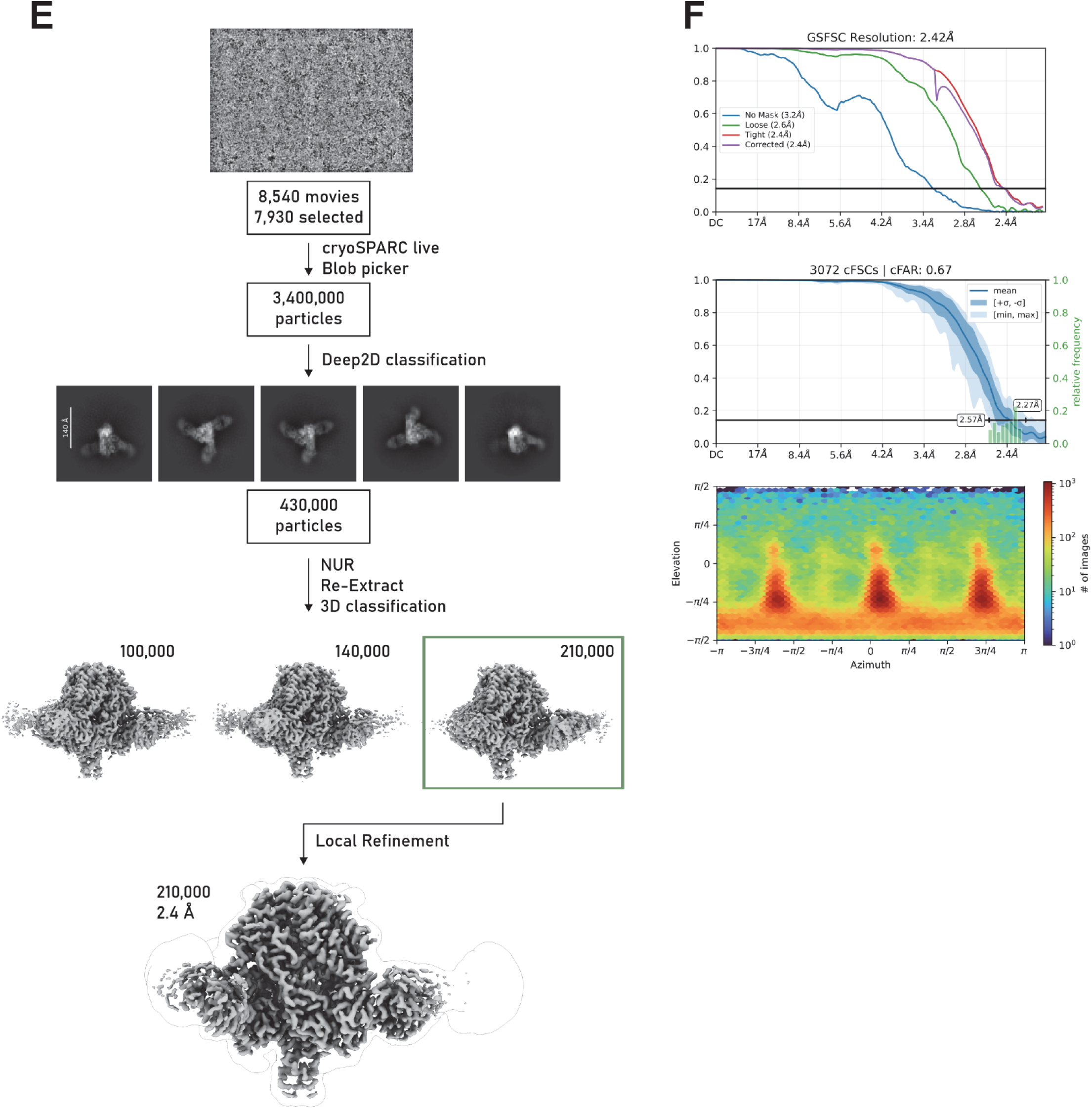
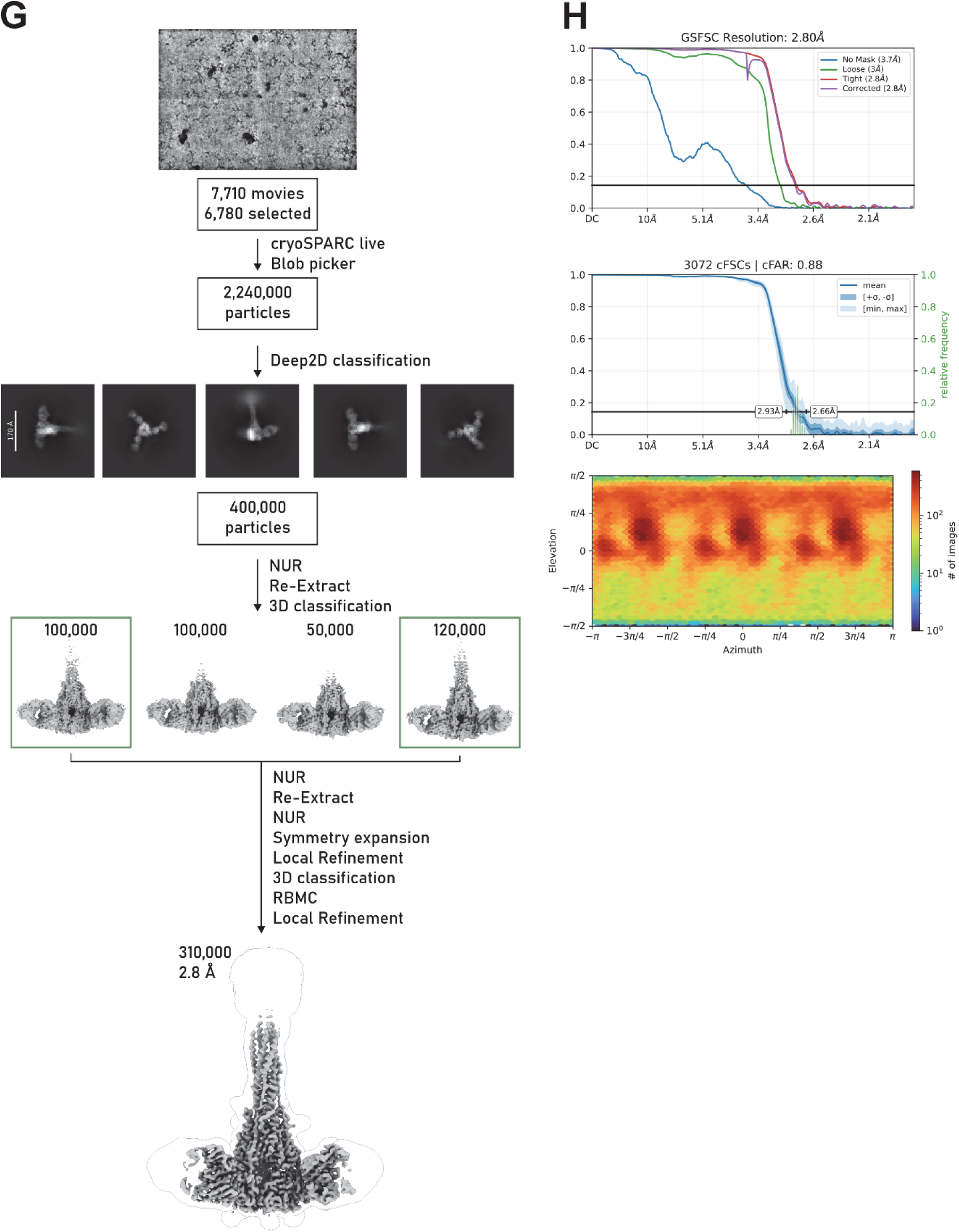
(A -H). Cryo-EM processing pipeline and map statistics. A-B: Cryo-EM processing pipeline (**A**) and final map statistics (**B**) for F_ECTO_-Y10F structure. **C-D**) Processing pipeline (**C**) and statistics (**D**) for F_ECTO_-Y10F-77.1 structure. **E-F**) Processing pipeline (**E**) and statistics (**F**) for F_ECTO_-H8 structure in prefusion conformation. **G-H**) Processing pipeline (**G**) and statistics (**H**) for F_ECTO_-H8 structure in postfusion conformation.

**Figure S2.**
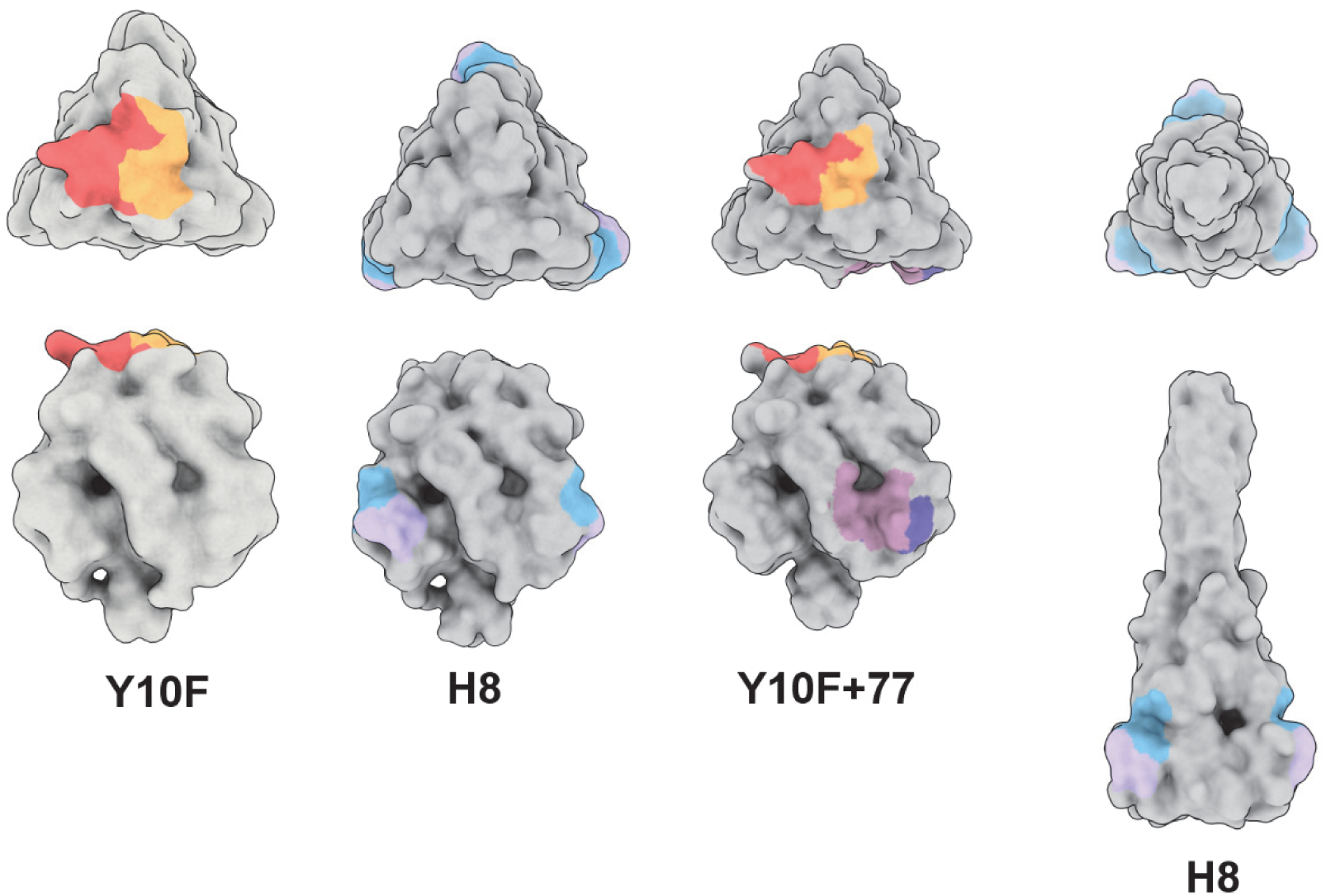
Antibody footprints of Y10F, H8, and 77.1 on the F_ECTO_ trimer in prefusion and postfusion conformations. Surface representations of the fusion ectodomain (rendered at 12 Å resolution) show the binding footprints of the antibodies characterized in this study. Y10F is shown with heavy and light chains in orange and yellow, respectively. H8 is shown with heavy and light chains in purple and blue. 77.1 is shown with heavy and light chains in pink and purple, respectively. The final panel displays the H8 epitope mapped onto the postfusion conformation of the F protein.

**Figure S3.**
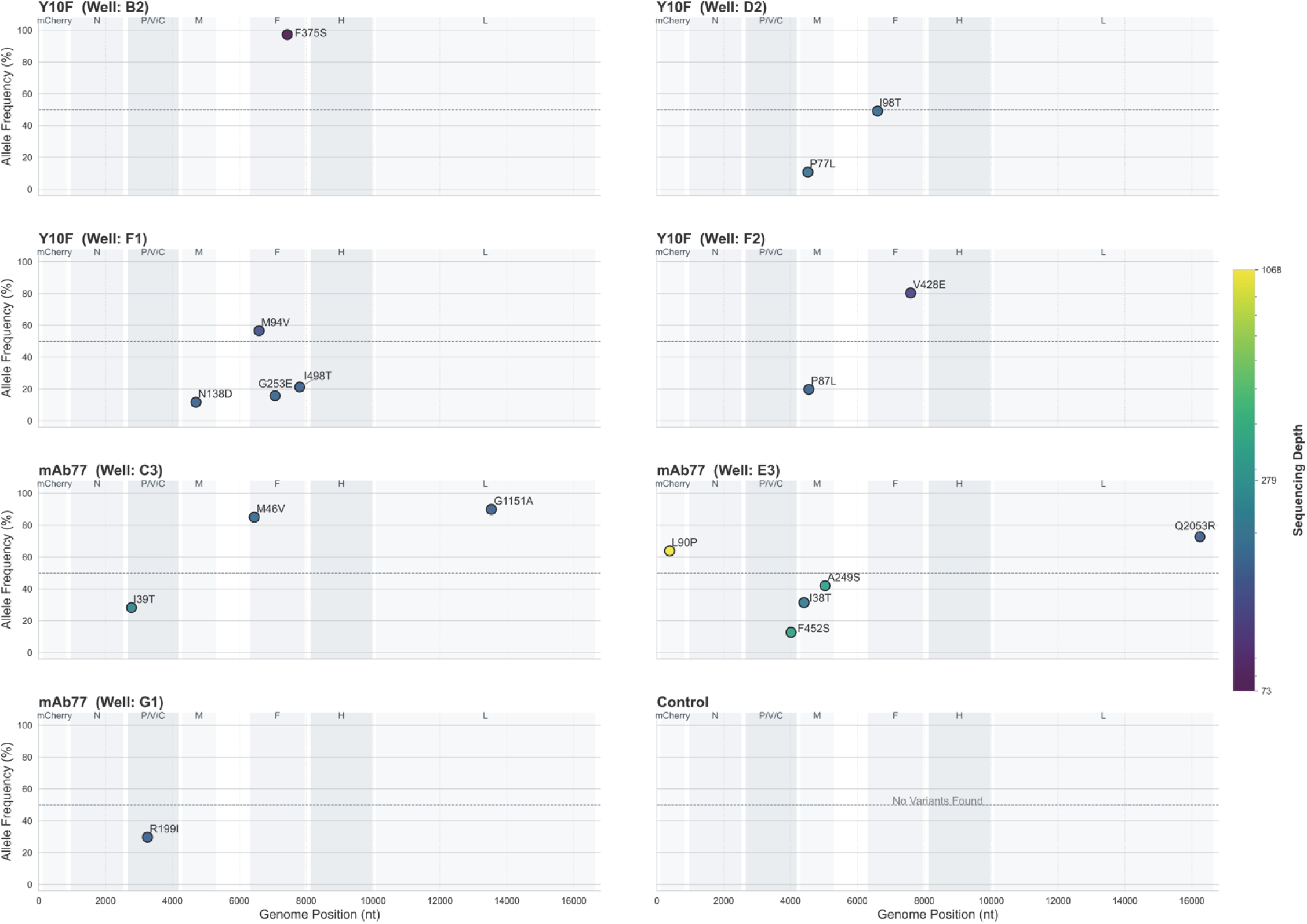
Allele frequency distribution of synonymous and nonsynonymous mutations in evolved viral populations. MeV was cultured in the presence of mAb 77, Y10F, or without any inhibitors for over 50 days as indicated in Materials and Methods. Nucleotide polymorphisms and corresponding amino acid changes relative to the wild-type strain are depicted. Refined mutational profile, filtered for allele frequencies >= 10% and sequencing depth >= 20. Points are color-coded to match the sequence depth, as indicated by the color bar at the bottom.

**Figure S4.**
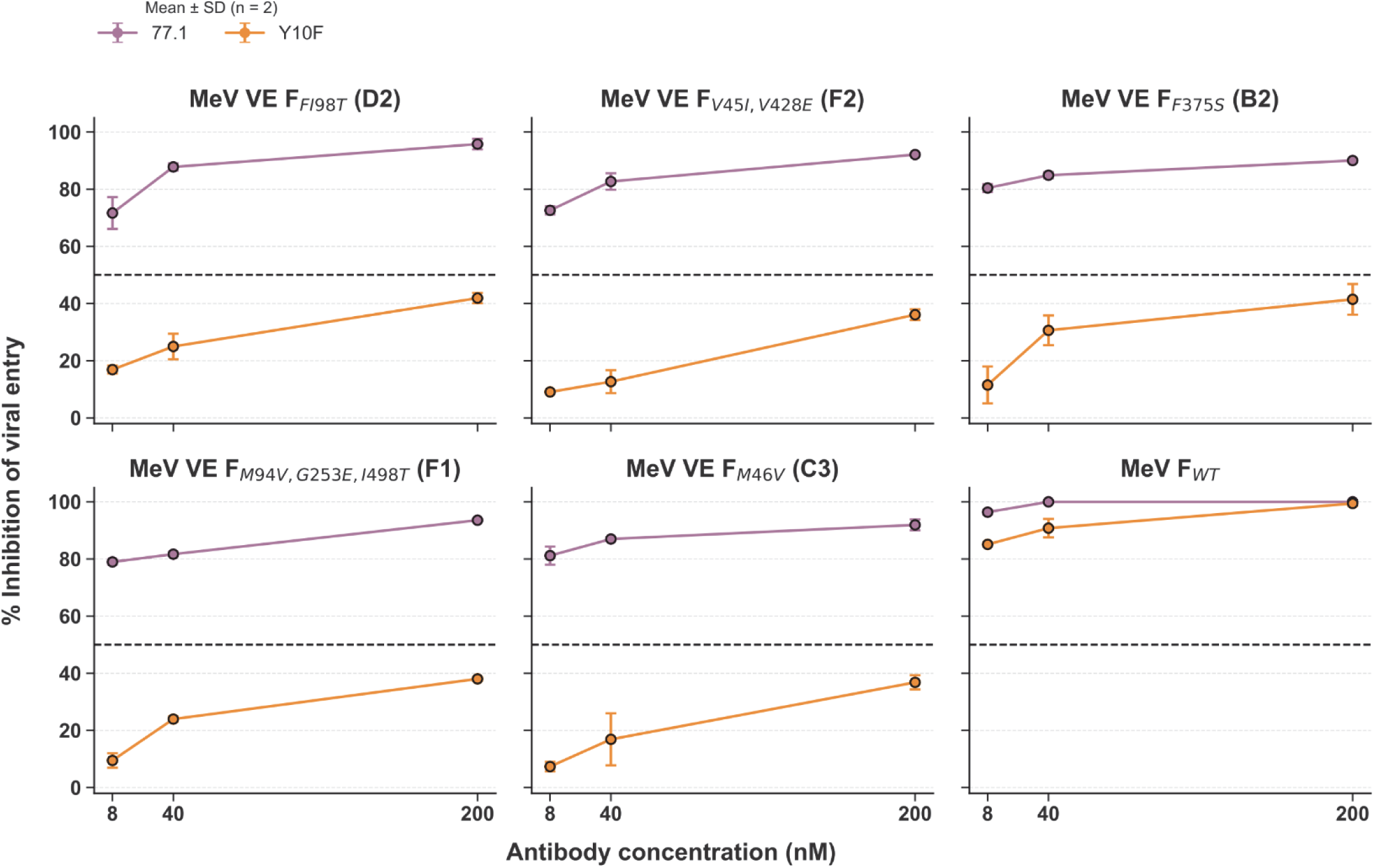
Inhibitory activity of viral populations following viral evolution. Viral entry inhibition by either mAb 77.1 (blue circles) or mAb Y10F (red squares) viral population after passages in the presence of Y10F (**A-D**) and 77.1 (**E**) compared to wt (**F**). 200pfu/well were incubated in the presence of the indicated concentration of the mAbs (x-axis). After 3 hours, the medium was replaced with a medium containing 2% methylcellulose Avicel and 2× complete medium (1:1). Two days post-infection, the number of infected cells was counted using a Cytation 5 (BioTek) and analyzed with Gen5 3.11 software. The percentage of inhibition was calculated relative to the no-treatment control. Data represent the mean ± SD of two independent experiments.

**Figure S5.**
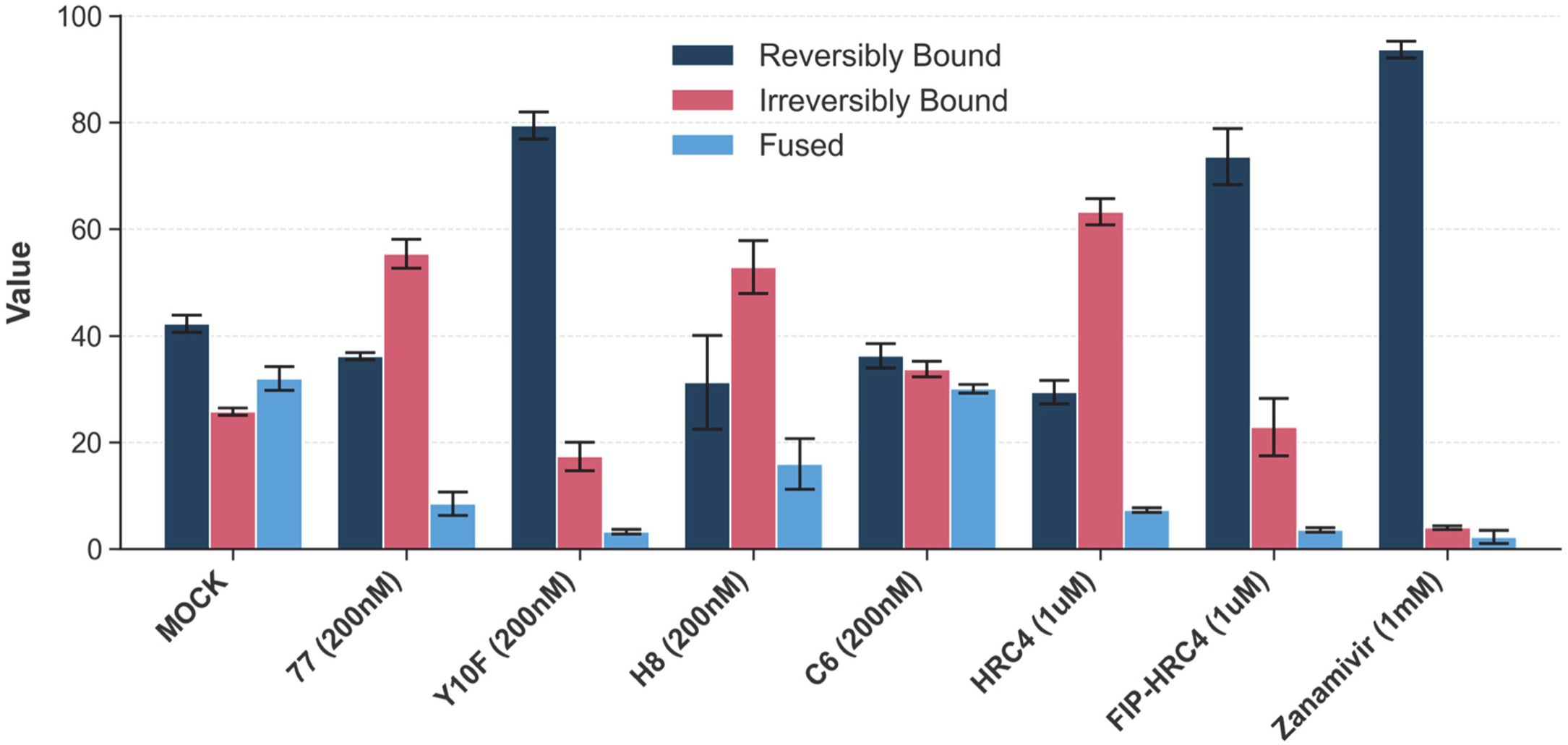
Detailed plot from Fig. 5C showing red blood cell assay incubation at 37°C for 60 min. Inhibitors and controls are shown, as are their concentrations.

**Table S1.**
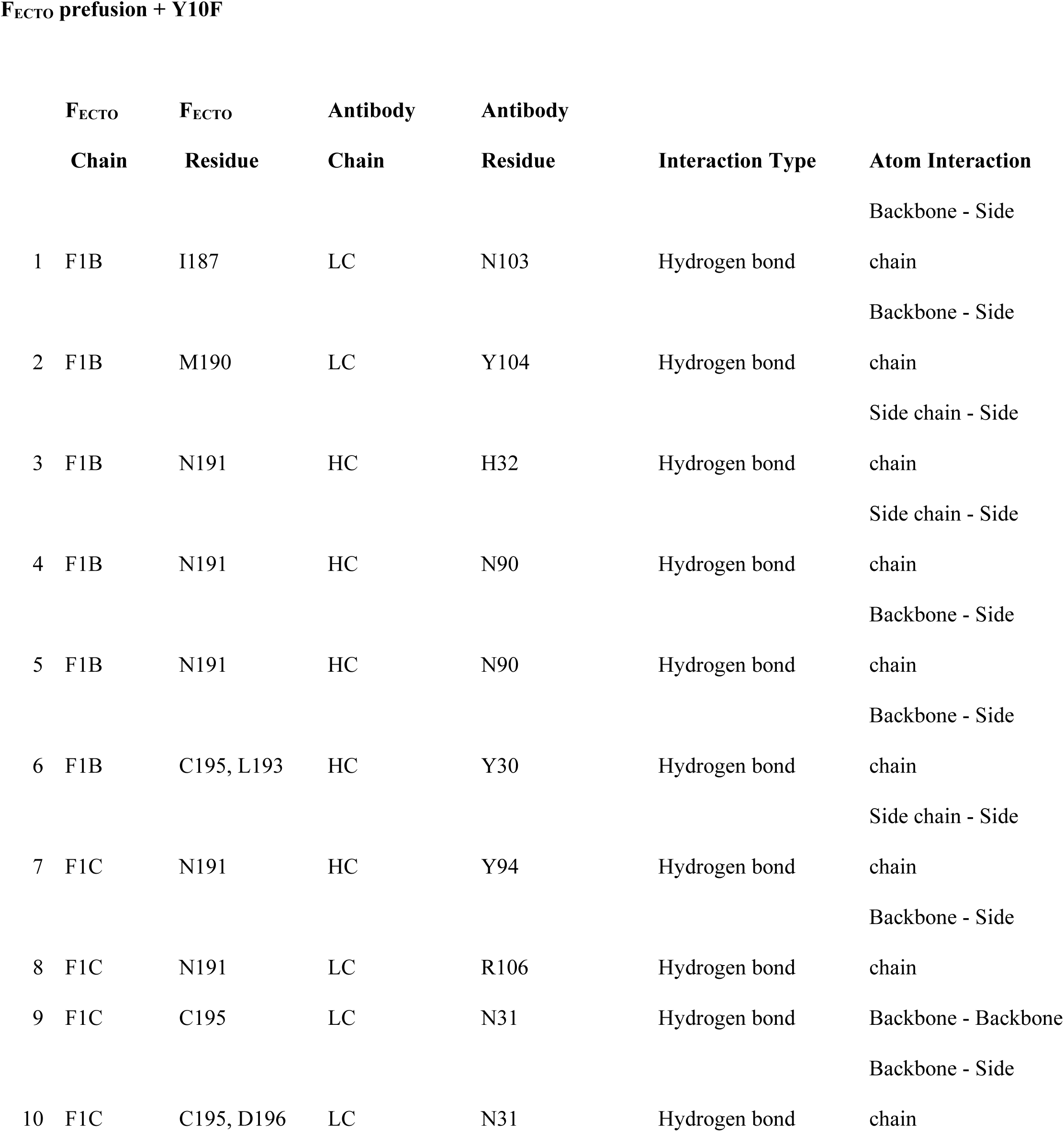
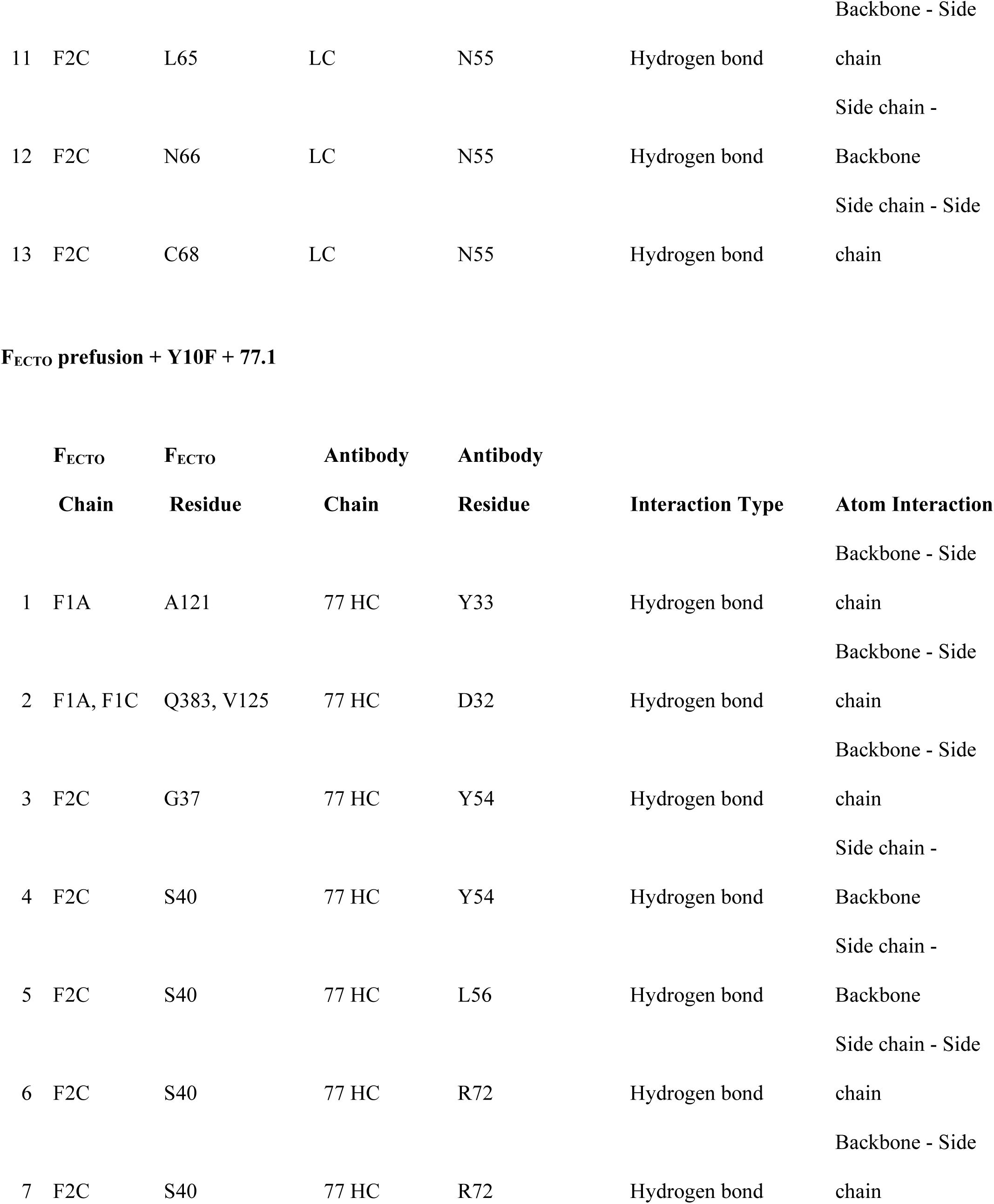
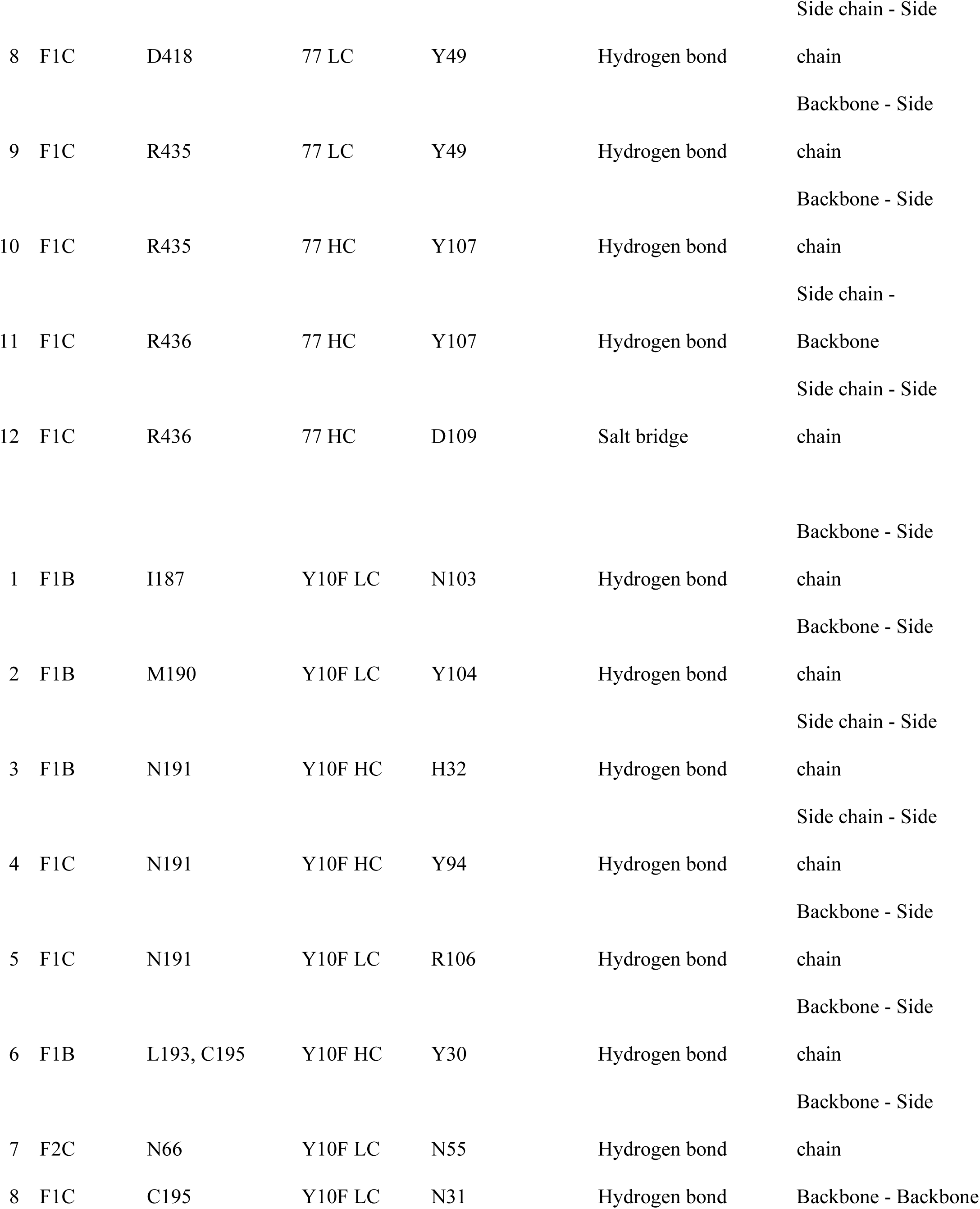
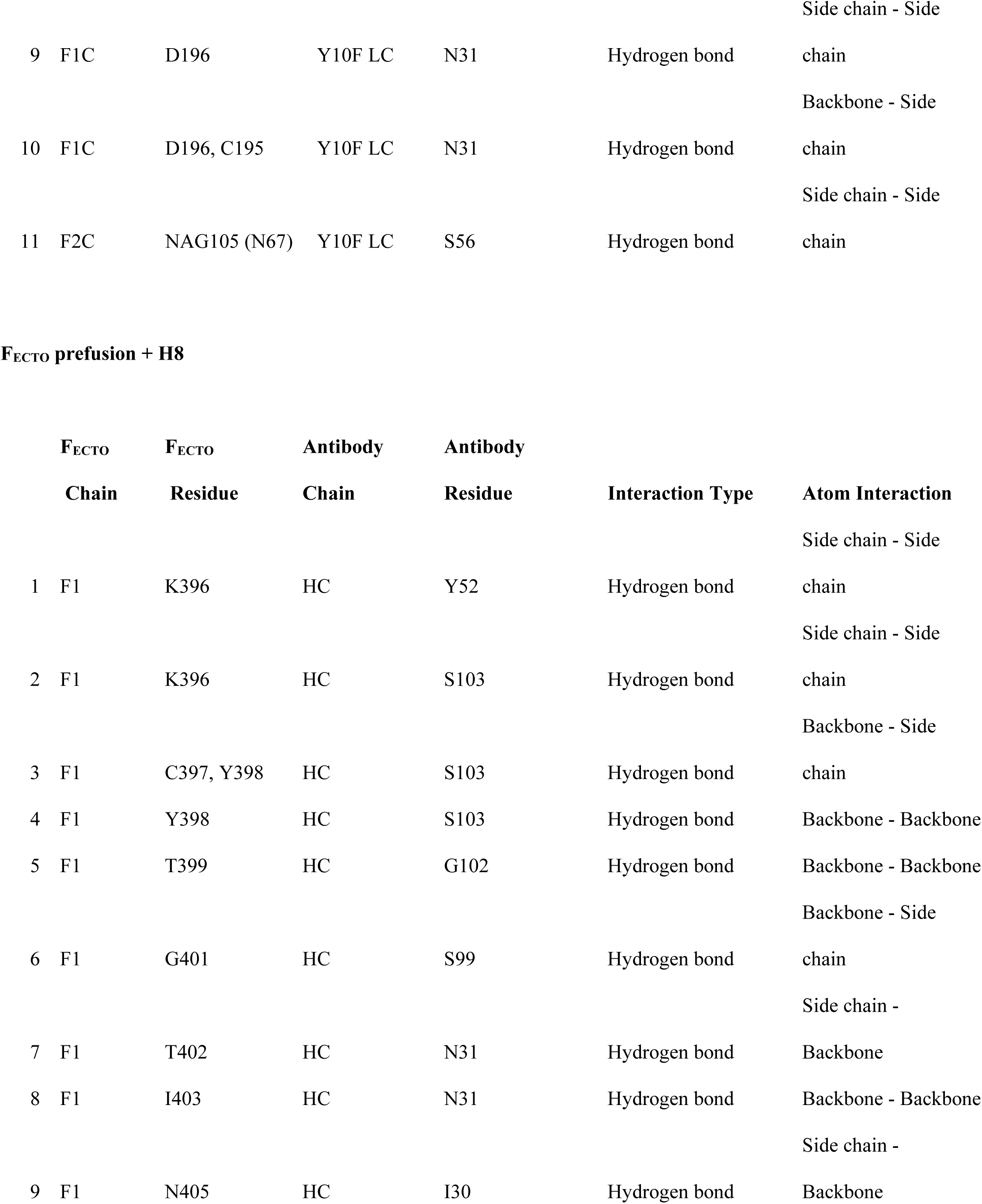
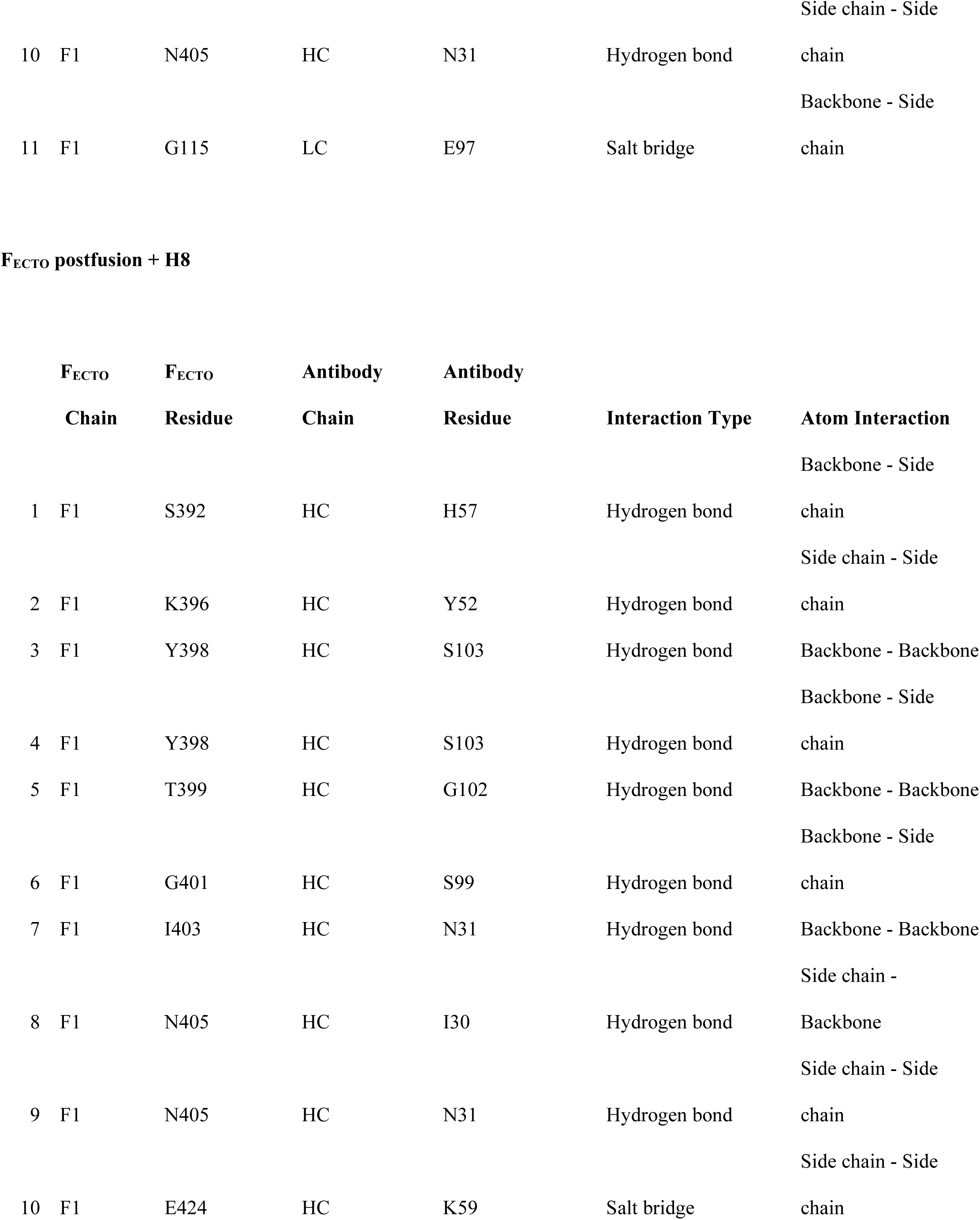
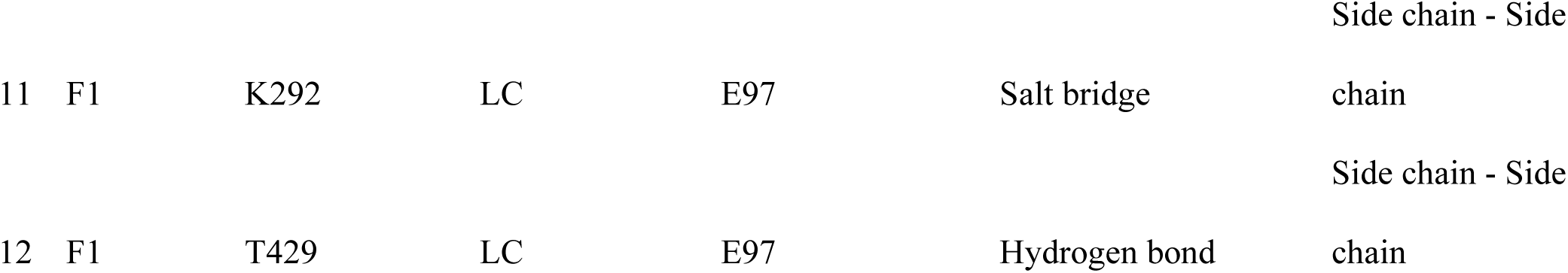
Summary F-mAb contact residues as determined by cryo-EM.

**Table S2.** Cryo-EM statistics from all recorded datasets.

|  | F<sub>ECTO</sub> + H8 (prefusion) | F<sub>ECTO</sub> + H8 (postfusion) | F<sub>ECTO</sub> + Y10F + 77 | F<sub>ECTO</sub> + Y10F |
| --- | --- | --- | --- | --- |
| Data Collection |  |  |  |  |
| Microscope and camera | Titan Krios G3 + Gatan K3 with Bio Quantum Energy Filter |  |  |  |
| Voltage (kV) |  | 300 kV |  |  |
| Magnification |  | 130,000 |  |  |
| Raw Pixel size (Å) |  | 0.66 |  |  |
| Electron dose (e<sup>-</sup>/Å<sup>2</sup>) |  | 70 |  |  |
| Energy Filter slit (eV) |  | 20 |  |  |
| Refinement |  |  |  |  |
| Final box size (px) | 320 | 584 | 380 | 464 |
| Final pixel size (Å) | 1.056 | 0.88 | 1.0768 | 0.88 |
| Final number of particles | 210,483 | 312,278 | 221,894 | 129,970 |
| Symmetry applied | C3 (relaxed) | C3 (relaxed) | C1 | C1 |
| Map and Model Statistics |  |  |  |  |
| 0.143 FSC Resolution (Å) | 2.42 | 2.8 | 2.82 | 2.57 |
| CC (mask) | 0.77 | 0.87 | 0.8 | 0.85 |
| 0.143 Masked Map/Model FSC (Å) | 2.4/2.4/2.7 | 2.7/2.8/3.0 | 2.7/2.8/3.1 | 2.5/2.5/2.8 |
| Model Composition |  |  |  |  |
| Chains | 14 | 15 | 13 | 12 |
| Protein Residues | 2024 | 1919 | 1811 | 1585 |

|  | **F<sub>ECTO</sub> + H8 (prefusion)** | **F<sub>ECTO</sub> + H8 (postfusion)** | **F<sub>ECTO</sub> + Y10F + 77** | **F<sub>ECTO</sub> + Y10F** |
| --- | --- | --- | --- | --- |
| Ligands | NAG: 15, BMA: 3, MAN: 4 | NAG: 9 | NAG: 16 | NAG: 15, BMA: 3, MAN: 4 |
| Water molecules | 140 | 0 | 0 | 63 |
| Total Atoms | 15835 | 14828 | 14059 | 12394 |
| **Model Quality** |  |  |  |  |
| MolProbity score | 1.5 | 1.33 | 1.22 | 1.36 |
| Clash score | 5.77 | 3.34 | 2.91 | 5.53 |
| Rotamer outliers (%) | 0.0 | 0.06 | 0.0 | 0.0 |
| CaBLAM outliers (%) | 1.42 | 1.66 | 1.13 | 1.29 |
| Ramachandran Plot (%) |  |  |  |  |
| Favored | 96.9 | 96.72 | 97.26 | 97.71 |
| Allowed | 3.1 | 3.28 | 2.74 | 2.29 |
| Outliers | 0.0 | 0.0 | 0.0 | 0.0 |

## References

1. Do, L.A.H. & Mulholland, K. Measles 2025. N Engl J Med (2025).

2. Anderer, S. Global Measles Cases Rose 20% in 1 Year as Vaccine Coverage Fell Short. JAMA 333, 279 (2025).

3. Anderer, S. Measles Cases in European Region Highest in More Than 25 Years. JAMA 333, 1658 (2025).

4. Anderer, S. US Measles Cases Hit 25-Year High. JAMA 334, 661 (2025).

5. Anderer, S. Measles Resurgence Puts People With HIV at Risk. JAMA (2025).

6. Stoneman, E.K. Measles. JAMA (2025).

7. Williamson, K.M., et al. A Cross-Sectional Study of Measles-Specific Antibody Levels in Australian Blood Donors-Implications for Measles Post-Elimination Countries. Vaccines (Basel) 12(2024).

8. Gidding, H.F., Quinn, H.E., Hueston, L., Dwyer, D.E. & McIntyre, P.B. Declining measles antibodies in the era of elimination: Australia’s experience. Vaccine 36, 507–513 (2018).

9. Modrof, J., et al. Measles Virus Neutralizing Antibodies in Intravenous Immunoglobulins: Is an Increase by Revaccination of Plasma Donors Possible? J Infect Dis 216, 977–980 (2017).

10. Lackner, C., Karbiener, M., Faltner, L., Farcet, M.R. & Kreil, T.R. Feasibility of identifying plasma donors with high measles neutralizing antibody concentrations for the use of producing a measles hyperimmune globulin for postexposure prophylaxis. Immunol Res 70, 365–370 (2022).

11. Anichini, G., et al. Seroprevalence to Measles Virus after Vaccination or Natural Infection in an Adult Population, in Italy. Vaccines (Basel) 8(2020).

12. Bolotin, S., et al. In Elimination Settings, Measles Antibodies Wane After Vaccination but Not After Infection: A Systematic Review and Meta-Analysis. J Infect Dis 226, 1127–1139 (2022).

13. Osman, S., et al. Population immunity to measles in Canada using Canadian Health Measures survey data - A Canadian Immunization Research Network (CIRN) study. Vaccine 40, 3228–3235 (2022).

14. Audet, S., et al. Measles-virus-neutralizing antibodies in intravenous immunoglobulins. J Infect Dis 194, 781–789 (2006).

15. Xiong, Y., et al. Age-related changes in serological susceptibility patterns to measles: results from a seroepidemiological study in Dongguan, China. Hum Vaccin Immunother 10, 1097–1003 (2014).

16. Pantaleo, G., Correia, B., Fenwick, C., Joo, V.S. & Perez, L. Antibodies to combat viral infections: development strategies and progress. Nat Rev Drug Discov 21, 676–696 (2022).

17. Pecenka, C., et al. Respiratory syncytial virus vaccination and immunoprophylaxis: realising the potential for protection of young children. Lancet 404, 1157–1170 (2024).

18. Corti, D., Purcell, L.A., Snell, G. & Veesler, D. Tackling COVID-19 with neutralizing monoclonal antibodies. Cell 184, 3086–3108 (2021).

19. de Swart, R.L., Yuksel, S. & Osterhaus, A.D. Relative contributions of measles virus hemagglutinin- and fusion protein-specific serum antibodies to virus neutralization. J Virol 79, 11547–11551 (2005).

20. de Swart, R.L., Yuksel, S., Langerijs, C.N., Muller, C.P. & Osterhaus, A. Depletion of measles virus glycoprotein-specific antibodies from human sera reveals genotype-specific neutralizing antibodies. J.Gen.Virol. 90, 2982–2989 (2009).

21. Harrison, S.C. Viral membrane fusion. Nature structural & molecular biology 15, 690–698 (2008).

22. White, J.M., Ward, A.E., Odongo, L. & Tamm, L.K. Viral Membrane Fusion: A Dance Between Proteins and Lipids. Annu Rev Virol 10, 139–161 (2023).

23. Duprex, W.P. & Dutch, R.E. Paramyxoviruses: Pathogenesis, Vaccines, Antivirals, and Prototypes for Pandemic Preparedness. J Infect Dis 228, S390–S397 (2023).

24. Rey, F.A. & Lok, S.M. Common Features of Enveloped Viruses and Implications for Immunogen Design for Next-Generation Vaccines. Cell 172, 1319–1334 (2018).

25. Jurgens, E.M., et al. Measles fusion machinery is dysregulated in neuropathogenic variants. mBio 6(2015).

26. Mathieu, C., et al. Measles Virus Bearing Measles Inclusion Body Encephalitis-Derived Fusion Protein Is Pathogenic after Infection via the Respiratory Route. J Virol 93, e01862–01818 (2019).

27. Zyla, D.S., et al. A neutralizing antibody prevents postfusion transition of measles virus fusion protein. Science 384, eadm8693 (2024).

28. Mathieu, C., et al. Molecular Features of the Measles Virus Viral Fusion Complex That Favor Infection and Spread in the Brain. mBio 12, e0079921 (2021).

29. Niewiesk, S. Current animal models: cotton rat animal model. Curr Top Microbiol Immunol 330, 89–110 (2009).

30. Bovier, F.T., et al. Inhibition of Measles Viral Fusion Is Enhanced by Targeting Multiple Domains of the Fusion Protein. ACS Nano (2021).

31. Stokke, J.L., et al. MMR Vaccine-Associated Disseminated Measles in an Immunocompromised Adolescent. N Engl J Med 385, 1246–1248 (2021).

32. Ferrari, M.J. & Moss, W.J. Rubella and measles: The beginning of the endgame. Science 388, 32–34 (2025).

33. Patel, M.K., et al. Progress Toward Regional Measles Elimination - Worldwide, 2000-2018. MMWR Morb Mortal Wkly Rep 68, 1105–1111 (2019).

34. Reich, A., Erlwein, O., Niewiesk, S., ter Meulen, V. & Liebert, U.G. CD4+ T cells control measles virus infection of the central nervous system. Immunology 76, 185–191 (1992).

35. Paules, C.I., Marston, H.D. & Fauci, A.S. Measles in 2019 - Going Backward. N Engl J Med 380, 2185–2187 (2019).

36. Mina, M.J., et al. Measles virus infection diminishes preexisting antibodies that offer protection from other pathogens. Science 366, 599–606 (2019).

37. Munoz-Alia, M.A., et al. Measles Virus Hemagglutinin epitopes immunogenic in natural infection and vaccination are targeted by broad or genotype-specific neutralizing monoclonal antibodies. Virus Res 236, 30–43 (2017).

38. Vos, R.A., et al. Seroepidemiology of Measles, Mumps and Rubella on Bonaire, St. Eustatius and Saba: The First Population-Based Serosurveillance Study in Caribbean Netherlands. Vaccines (Basel) 7(2019).

39. van den Hof, S., Berbers, G.A., de Melker, H.E. & Conyn-van Spaendonck, M.A. Sero-epidemiology of measles antibodies in the Netherlands, a cross-sectional study in a national sample and in communities with low vaccine coverage. Vaccine 18, 931–940 (1999).

40. Mathieu, C., et al. Single-chain variable fragment antibody constructs neutralize measles virus infection in vitro and in vivo. Cell Mol Immunol 18, 1835–1837 (2021).

41. Malvoisin, E. & Wild, F. Contribution of measles virus fusion protein in protective immunity: anti-F monoclonal antibodies neutralize virus infectivity and protect mice against challenge. J Virol 64, 5160–5162 (1990).

42. Byrne, P.O., et al. Structural basis for antibody recognition of vulnerable epitopes on Nipah virus F protein. Nat Commun 14, 1494 (2023).

43. Angius, F., et al. Analysis of a Subacute Sclerosing Panencephalitis Genotype B3 Virus from the 2009-2010 South African Measles Epidemic Shows That Hyperfusogenic F Proteins Contribute to Measles Virus Infection in the Brain. J Virol 93, e01700–01718 (2019).

44. Meng, E.C., et al. UCSF ChimeraX: Tools for structure building and analysis. Protein Sci 32, e4792 (2023).

45. Pettersen, E.F., et al. UCSF ChimeraX: Structure visualization for researchers, educators, and developers. Protein Sci 30, 70–82 (2021).

46. Emsley, P. & Cowtan, K. Coot: model-building tools for molecular graphics. Acta Crystallogr D Biol Crystallogr 60, 2126–2132 (2004).

47. Liebschner, D., et al. Macromolecular structure determination using X-rays, neutrons and electrons: recent developments in Phenix. Acta Crystallogr D Struct Biol 75, 861–877 (2019).

48. Agirre, J., et al. The CCP4 suite: integrative software for macromolecular crystallography. Acta Crystallogr D Struct Biol 79, 449–461 (2023).

49. Greninger, A.L., et al. Rapid Metagenomic Next-Generation Sequencing during an Investigation of Hospital-Acquired Human Parainfluenza Virus 3 Infections. J Clin Microbiol 55, 177–182 (2017).

50. Di Tommaso, P., et al. Nextflow enables reproducible computational workflows. Nat Biotechnol 35, 316–319 (2017).

